# Surprising Phenotypic Diversity of Cancer-associated mutations of Gly 34 in the Histone H3 tail

**DOI:** 10.1101/2020.12.10.419184

**Authors:** Brandon R. Lowe, Rajesh K. Yadav, Ryan A. Henry, Patrick Schreiner, Atsushi Matsuda, Alfonso G. Fernandez, David Finkelstein, Margaret Campbell, Satish Kallappagoudar, Carolyn M. Jablonowski, Andrew J. Andrews, Yasushi Hiraoka, Janet F. Partridge

## Abstract

Sequencing of cancer genomes has identified recurrent somatic mutations in histones, termed oncohistones, which are frequently poorly understood. Previously we showed that fission yeast expressing only the H3.3G34R mutant identified in aggressive pediatric glioma had reduced H3K36 trimethylation and acetylation, increased genomic instability and replicative stress, and defective homology-dependent DNA damage repair (Yadav et al., 2017). Here we show that surprisingly distinct phenotypes result from G34V (also in glioma) and G34W (giant cell tumors of bone) mutations, differentially affecting H3K36 modifications, subtelomeric silencing, genomic stability, sensitivity to irradiation, alkylating agents, hydroxyurea and influencing DNA repair. In cancer, only one of thirty alleles encoding H3 is mutated. Whilst co-expression of wild-type H3 rescues most G34 mutant phenotypes, G34R causes dominant hydroxyurea sensitivity and homologous recombination defects, and dominant subtelomeric silencing. Together, these studies demonstrate the complexity associated with different substitutions at even a single residue in H3 and highlight the utility of genetically tractable systems for their analysis.

## Introduction

The fundamental regulation of DNA within eukaryotes is coordinated through highly conserved histone proteins that package DNA into the nucleus. Proteins that regulate post-translational modifications of histone proteins or of DNA, or that control nucleosome assembly, disassembly or movement, are frequently targeted in cancer (Huether et al., 2014; Roy, Walsh, & Chan, 2014; Shen & Laird, 2013). More recently, mutations within the histone genes themselves have been identified in disease (Behjati et al., 2013; Lu et al., 2016; Nacev et al., 2019; Schwartzentruber et al., 2012; Tessadori et al., 2017; Wu et al., 2012). These histone mutations arise predominantly in one of two genes encoding the histone variant protein H3.3. H3.3 is expressed throughout the cell cycle and, in contrast to the replication-dependent histone H3.1 and H3.2, is deposited into chromatin outside of S-phase (reviewed in (Kallappagoudar et al., 2015)). Oncogenic H3.3 mutations frequently alter residues that are key sites of post-translational modification (K27M or K36M), or mutate G34, which affects the methylation of the neighboring K36 (H3K36me (Lewis et al., 2013)). The K27M and K36M mutants, found in 84% of diffuse intrinsic pontine gliomas and 95% chondroblastomas respectively (Behjati et al., 2013; Schwartzentruber et al., 2012; Wu et al., 2012), exert dominant effects by inhibition of the relevant methyltransferase complexes through binding the mutant histone tails, reducing methylation of total cellular histone H3 pools at K27 or K36 respectively (Brown et al., 2014; Fang et al., 2016; Jain et al., 2019; Lewis et al., 2013; Lu et al., 2016; Zhang et al., 2017). We know much less, however, about the role of histone mutants that lack obvious dominant effects on post-translational modification of the remaining histone pools (Bjerke et al., 2013; Lewis et al., 2013; Nacev et al., 2019; Tessadori et al., 2017). For example, although H4 K91 mutants (Q and R) impact acetylation and ubiquitination of the mutant H4 tail, and H3.3 G34R and V decrease K36me3 on the mutant H3.3 tail, no alterations in post-translational modification have been reported on the wild type H4 or H3 proteins expressed in these cells (Lewis et al., 2013; Tessadori et al., 2017). However, pronounced redistribution of the K36me3 mark was seen in a pediatric glioblastoma cell line derived from an H3.3 G34V patient (KNS42), correlating with a switch to transcriptional activation of genes normally expressed in early embryonic development (Bjerke et al., 2013). Whether this change is caused by the histone mutation, is a downstream effect of transcriptional change, or the result of another mutation frequently found in H3.3 G34V mutant glioma, for example mutation of p53 or H3.3 chaperone and remodeler proteins DAXX and ATRX (Bjerke et al., 2013; Korshunov et al., 2016; Schwartzentruber et al., 2012; Wu et al., 2012), is currently unclear.

We set out to address the role of mutations in histone H3, one of the most highly conserved proteins in eukaryotes, using the highly tractable fission yeast. This system excels for evaluation of the biological effects of mutations divorced from confounding effects of additional mutations in tumors, and can be stripped of the complexity of additional wild-type histone H3 variants within cells. We previously reported that fission yeast manipulated to express only G34R mutant histone H3 (H3-G34R) showed defective H3K36 acetylation (H3K36ac), and reduced levels of H3K36me3 (Yadav et al., 2017). These chromatin changes correlated with some transcriptional change including upregulation of antisense transcripts and enhanced subtelomeric silencing, as well as defects in replication and homologous recombination (HR) which we hypothesized were causal of genomic instability in these cells. However, it remained unclear if particular chromatin changes could be linked to specific defects.

Here, we investigate other mutants of H3G34. H3-G34V mutation is less frequent than G34R in pediatric high grade glioma, but the tumors occur in the same cortical location in patients of similar age (Schwartzentruber et al., 2012; Wu et al., 2012). Based on these findings, we expected similar behavior of H3-G34V and H3-G34R mutants. Surprisingly, although H3K36me3 was reduced in both mutants, other phenotypes were quite distinct, with G34V showing no defect in H3K36 acetylation, no chromosome loss, no replicative stress and no defects in HR, but enhanced sensitivity to gamma (γ) irradiation. We queried whether these differences were due to the size or the charge of the substitution. By generating a panel of H3-G34 mutant yeast, we found that G34K, which like G34R incorporates a basic charge, also showed reduced K36me3 and chromosome loss but G34W, a bulky uncharged substitution found in H3.3 in nearly all giant cell tumors of bone (Behjati et al., 2013), did not alter H3K36me3 or genome stability and did not sensitize cells to the DNA damaging agents tested. We further examined which effects were dominant, and found that only some, including the HU sensitivity and HR defects in H3-G34R mutants, and silencing of subtelomeric domains in both H3-G34R and G34K mutants were evident in cells expressing a mixture of wild type and mutant H3. In contrast, chromosome segregation was corrected and most DNA damage sensitivities were lost when cells additionally express wild type H3. This study reveals the pivotal role of H3 Gly 34 in orchestrating many cellular responses, some that apparently act on a local level and others able to exert dominant effects.

## Results

Fission yeast have three genes that code for a single histone H3 protein (Fig. 1a) (Matsumoto & Yanagida, 1985). Strains were derived that express only one H3 and one H4 gene (*hht2*^*+*^ and *hhf2^+^*) (Mellone et al., 2003), which maintain histone protein levels similar to those in wild-type strains (Yadav et al., 2017). Previously, we introduced a mutation into *hht2*^*+*^ (Fig. 1a) to generate strains that express only the G34R mutant form of histone H3 (Yadav et al., 2017). In the current study, we derived a panel of strains that express only H3-G34 mutants, and compared them with H3-G34R strains. “Single copy” histone H3 and H4 strains were used throughout and are named **H3-WT** and **H3-G34X** in the text, and “**WT**” and “**G34X**” in the figures. Additional mutations were also generated in the single H3 background unless denoted (**3xH3**) representing strains with 3 copies of H3/H4 (Supplementary Table S1).

**Figure 1.**
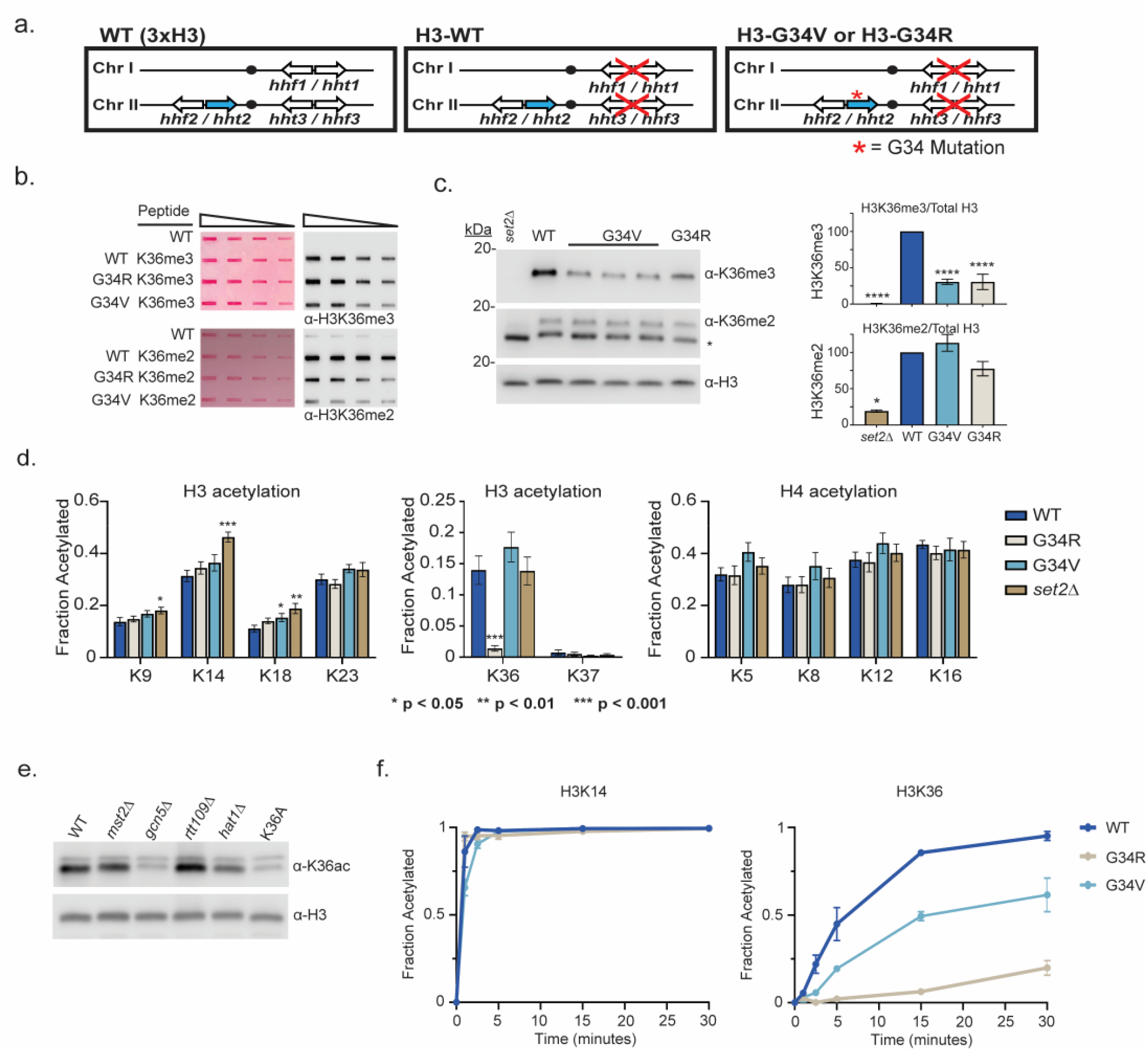
Differential modification of H3K36 in H3-G34R and H3-G34V mutants. (a) Scheme of the histone H3 (*hht*) and histone H4 (*hhf*) genes in *S. pombe* highlighting the H3 gene (*hht2*) in which mutations were engineered (blue). (b) Dot blot analysis to quantitatively assess recognition of WT, G34R and G34V peptides bearing K36 di or tri-methyl modifications by anti-K36 methyl antibodies. Ponceau stained blots were used as the loading control (left). (c) Western blot analysis of K36me2, K36me3, and total H3 in H3-WT, H3-G34V, H3-G34R and *set2Δ* chromatin extracts. Star marks nonspecific band. (Right) quantification of K36 methylation relative to total H3 (K36me3: 3 replicates for H3-WT, *set2Δ* and H3-G34R, and 8 replicates for H3-G34V; K36me2: 2 replicates for H3-WT, H3-G34R and *set2Δ* and 5 replicates for H3-G34V). For K36me3 blot, **** represents significance difference p<0.0001 from H3-WT strain. (d) Mass spectrometry-based quantification of acetylation of specific lysines in histone H4 and H3 tails in histones purified from H3-WT, H3-G34R, H3-G34V and *set2Δ* strains. H3-WT, H3-G34R and *set2Δ* data (9 biological replicates, and H3-G34V (6 biological replicates). H3K27ac analysis was excluded as highly variable. (e) Western blot analysis of H3K36ac in lysates of (3xH3) WT, and acetyltransferase mutants *mst2Δ, gcn5Δ, hat1Δ,* and *rtt109Δ* with sole copy H3-K36A negative control (anti-H3K36ac Abnova PAB31320), and total H3 as loading control. (f) *In vitro* histone acetylation assay using recombinant Gcn5 and recombinant WT, G34R, or G34V H3 and monitoring H3K14ac and H3K36ac. Data from each time point represents the mean +/− SEM from 3 biological replicates.

### H3K36me3 is reduced in H3-G34V mutants

In fission yeast, all stages of H3 K36 methylation (mono, di and tri) are carried out by a single enzyme, Set2 (Morris et al., 2005). Experiments monitoring H3K36 methylation by mass spectrometry in human cells have shown that expression of G34R or V mutant H3.3 reduces H3K36me2 and me3 on the same histone tail (Lewis et al., 2013). Since H3-G34R mutant fission yeast exhibit a pronounced reduction in H3K36me3 (Yadav et al., 2017), we asked whether H3K36me was similarly affected by H3-G34V. We first characterized antibodies for recognition of H3K36 methylation in peptides containing a H3G34V mutant tail (Fig 1b, Supplementary Table S2). Using an anti-K36me3 antibody that bound efficiently to G34V mutant tails, western analyses showed a marked reduction in H3K36me3 in chromatin extracts from H3-G34V compared to H3-WT cells (Fig. 1c). In contrast, H3K36me2 appeared unchanged on western analysis of H3-G34V compared to H3-WT strains, but is upregulated in H3-G34V cells since antibody recognition of K36me2 is notably reduced on the G34V mutant H3 tail. A similar effect on K36me2 was seen in G34R cells (Yadav et al., 2017). Thus H3K36me3 is reduced and H3K36me2 accumulates in both H3-G34V and H3-G34R cells. Since Set2 is the sole K36 methyltransferase in fission yeast, this result suggests that there may be a defect in Set2-mediated trimethylation of K36me2 G34V/R templates, or that there is heightened activity of a K36me3 demethylase. To probe these possibilities, we performed ChIP analysis of Set2 in G34V/R cells, but saw no change in Set2 association with chromatin at the sites tested (Fig. 1-supplementary fig. 1a). Second, we asked whether loss of Epe1, a proposed H3 demethylase (Trewick, Minc, Antonelli, Urano, & Allshire, 2007), influences levels of H3K36 methylation, but we saw no consistent change in K36me2 or me3 in *epe1Δ* strains (Fig. 1-supplementary fig. 1b).

### H3-G34R but not H3-G34V cells exhibit differences in H3 acetylation

Since H3-G34R mutation results in a marked reduction in H3K36ac (Yadav et al., 2017), we asked whether K36ac was also affected by G34V mutation. Targeted quantitative mass spectrometry of histone H3 and H4 acetylation was performed on histones extracted from H3-WT, H3-G34V, H3-G34R and *set2Δ* cells (Fig. 1d, Supplementary Table S3) (Kuo & Andrews, 2013). We found that H3K36ac was greatly reduced in H3-G34R, but was unaffected in H3-G34V cells. In conclusion, H3-G34R, but not G34V, appears to inhibit efficient acetylation of H3 K36, whereas both mutants show loss of H3K36me3.

### Gcn5 acetylates H3K36, but cannot acetylate K36 on H3-G34R tail in vitro

To determine which histone acetyltransferase mediates H3K36ac, we assessed H3K36ac levels in cells lacking various histone acetyltransferases. Gcn5 has previously been implicated in H3K36ac (Pai et al., 2014) and consistent with this, using antibody that was largely specific to K36ac (Fig. 1-supplementary fig. 1c), we found that *gcn5Δ* cells have low H3K36ac levels, similar to H3-K36A cells which lack H3K36ac (Fig. 1e). To extend this analysis, since GCN5 has many targets including H3K14, H3K18 and H3K23, we asked if the specific loss of K36ac on H3-G34R mutant H3 could be recapitulated in vitro. Using recombinant fission yeast Gcn5 with recombinant WT or G34 mutant histone H3s, we used mass spectroscopy to monitor acetylation at individual sites in the H3 tail over time (Kuo & Andrews, 2013; Kuo, Henry, & Andrews, 2014). Whereas Gcn5 acetylated most sites on WT, G34R and G34V mutant H3 templates equivalently, acetylation of K36 on G34R H3 was strongly reduced, and was slowed on G34V H3 (Fig. 1f and Fig. 1-supplementary fig. 1d). Together, the in vivo and in vitro data suggest that the G34R mutation specifically reduces acetylation on H3 K36, while acetylation at other sites remains largely unaffected. This data is consistent with structural studies of Gcn5-substrate complexes which predicted that residues just N-terminal to the reactive lysine would modulate substrate affinity, and that replacement of the small residue (glycine or alanine) at position −2 (in this case H3G34 relative to H3K36 target) with a larger residue would likely impact Gcn5 activity (Fig. 1-supplementary fig. 1e, (Poux & Marmorstein, 2003)).

### H3-G34V mutants are not sensitive to replication stress

Altered H3K36 post-translational modification is associated with defective repair of DNA damage (Jha & Strahl, 2014; Li et al., 2013; Pai et al., 2014; Pfister et al., 2014). Consistent with this, H3-G34R cells with reduced H3K36ac and H3K36me3 are sensitive to replicative stress (Yadav et al., 2017). To query the role of H3K36ac in replicative stress, we plated 5 fold serial dilutions of G34V and G34R cells onto media containing drugs, and assessed cell growth after several days (Fig. 2a). H3-G34R and *set2Δ* cells were sensitive to chronic exposure to hydroxyurea (HU, a ribonucleotide reductase inhibitor that depletes dNTPs), whereas H3-G34V cells were not. G34V cells were also not sensitive to DNA alkylation (methyl methanesulfonate, MMS) whereas both G34R and *set2Δ* cells were sensitive. Finally, H3-G34V cells were unaffected by exposure to the DNA topoisomerase I ligase inhibitor camptothecin (CPT), whereas H3-G34R and *set2Δ* showed slight or pronounced sensitivity respectively. At the concentrations used, these genotoxins predominantly affect DNA replication or require DNA replication to inflict damage. Thus while H3-G34R and *set2Δ* are sensitive to several forms of replicative stress, H3-G34V cells are not. Since both H3-G34V and H3-G34R have reduced K36me3, the sensitivity of H3-G34R cells to replicative stress cannot be attributed to a reduction in H3K36me3, but may be due to loss of H3K36ac.

**Figure 2.**
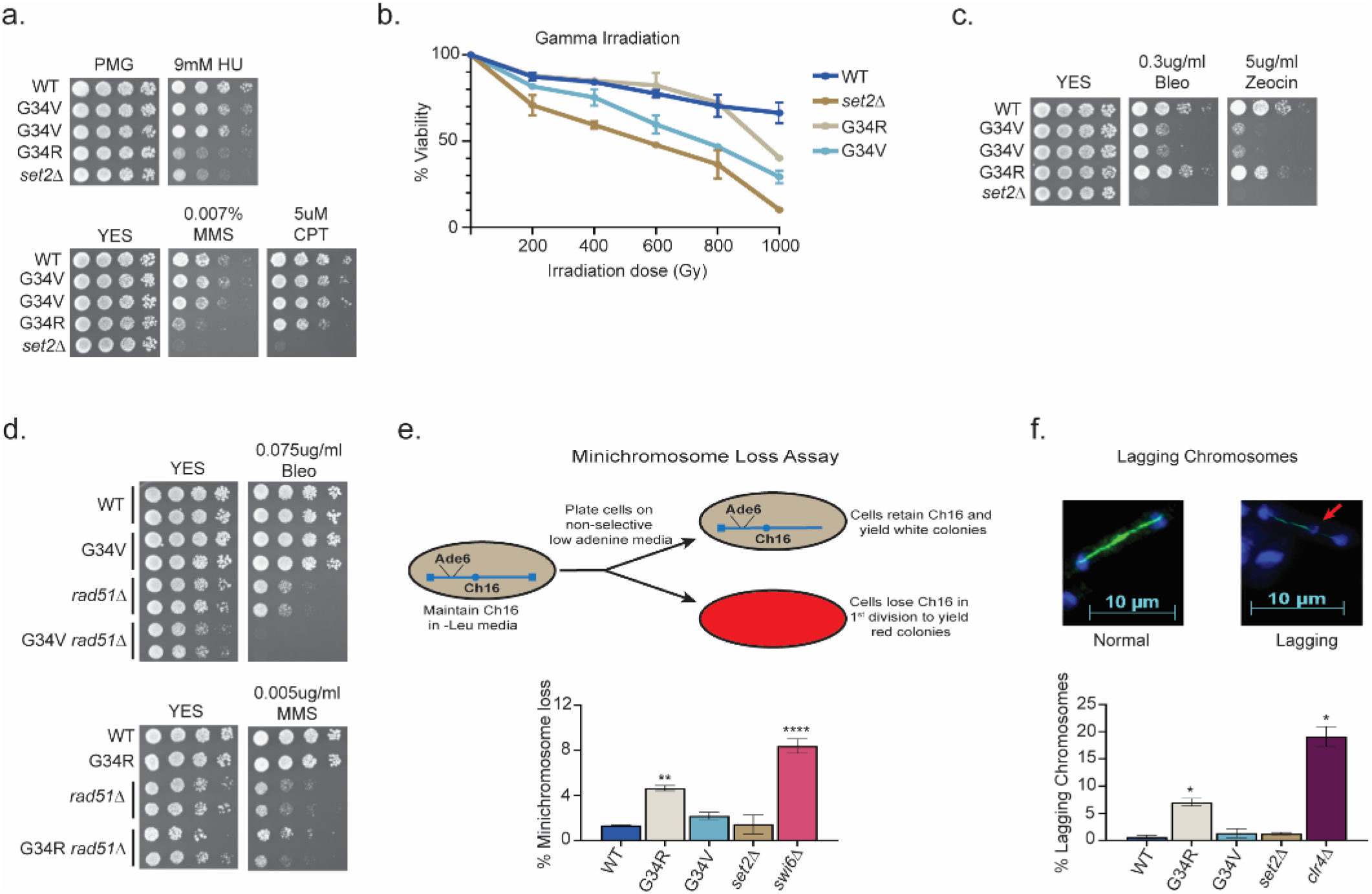
DNA damage sensitivity and chromosomal stability differ between H3-G34R and H3-G34V mutants. (a) 5-fold serial dilutions showing the effect of hydroxyurea (HU; a ribonucleotide reductase inhibitor that depletes dNTPs), methyl methanesulfonate (MMS; a DNA alkylating agent) and camptothecin (CPT; blocks topoisomerase 1 ligase activity) on growth of the indicated strains. (b) Effect of γ-irradiation (IR) exposure on viability of H3-WT, H3-G34R, H3-G34V, *set2Δ,* and H3-G34V *set2Δ* cells. Data represent mean +/− SEM from 2 independent experiments using 4 biological replicates. (c) Serial dilution assay showing the effect of bleomycin and zeocin, two irradiation mimetics, on the indicated strains. (d) Serial dilution assay assessing epistasis of mutants with HR pathway. Bleomycin sensitivity of H3-WT, H3-G34V, *rad51Δ,* and H3-G34V *rad51Δ* cells (top) and MMS sensitivity of H3-WT, H3-G34R, *rad51Δ,* and H3-G34R *rad51Δ* cells (bottom). (e) Frequency of cells that lose the non-essential minichromosome Ch16 in H3-WT, H3-G34V, H3-G34R, *set2Δ* and *swi6Δ* cells. Mean +/− SEM from 4-8 biological replicates is shown. ** denotes a significant difference of p<0.01 and **** p<0.0001 compared with the H3-WT strain. (f) Example of a normal anaphase and one with a lagging chromosome (red arrow) (top). Frequency of late anaphase cells with a lagging chromosome in H3-WT, G34R, G34V, *set2Δ,* and *clr4Δ* (bottom). Mean +/− SEM from 4-8 biological replicates. * represents significant difference of p<0.05 with the H3-WT strain.

### H3-G34V, but not H3-G34R mutants are sensitive to gamma irradiation and IR mimetics

Cells lacking *set2*^*+*^ are sensitive to γ-irradiation and irradiation mimetics (Pai et al., 2014) but H3-G34R cells are not (Yadav et al., 2017). We tested H3-G34V cells for sensitivity to γ-IR by transiently irradiating H3-WT, H3-G34R, H3-G34V, and *set2Δ* cells, and comparing viability by plating single cells and scoring colony formation after several days of growth (Fig. 2b). Notably, H3-G34V cells were sensitive to γ-irradiation, whereas as seen before, H3-G34R were not. H3-G34V and *set2Δ* cells were also sensitive to chronic exposure to the irradiation mimetics, bleomycin and zeocin, whereas H3-G34R were not (Fig. 2c). Thus the H3-G34V but not H3-G34R mutation renders cells sensitive to both γ-irradiation and to chronic exposure to irradiation mimetics.

### H3-G34V mutants are not defective for DNA break repair by homologous recombination

We next tested HR-mediated double-strand (ds) break repair efficiency in H3-G34V cells, since H3-G34R and *set2Δ* mutants are defective in HR-mediated DNA repair (Yadav et al., 2017). *leu1-32* mutant cells were transformed with a fragment of wild-type *leu1*^*+*^ and *leu1*^*+*^ transformants that arose by HR were scored (see Fig. 2-supplementary fig. 1a). In comparison to *set2Δ* and H3-G34R cells, which showed significant reduction in HR activity, the H3-G34V mutant showed no defect in HR (Fig. 2-supplementary fig. 1b). However, this assay probes HR efficiency at a single site, so we next performed a genetic epistasis analysis to ask more generally whether H3-G34V cells were competent for HR. To this end, we asked whether the DNA damage sensitivity of H3-G34V cells is enhanced on deletion of the major HR repair protein Rad51, which would suggest that HR is proficient in H3-G34V cells. Indeed we found that combination of *rad51Δ* and H3-G34V mutation enhanced DNA damage sensitivity, whereas combination of *rad51Δ* with H3-G34R rendered cells no more sensitive than *rad51Δ* (Fig. 2d). Together these assays indicate that H3-G34V cells rely on HR for efficient resolution of DNA damage, whereas H3-G34R cells are defective for HR. Note that we could not use epistasis analyses to probe other types of DNA damage repair in H3-G34V mutants since H3-G34V cells are only sensitive to IR or IR mimetics, and mutants defective in non-homologous end joining (NHEJ), such as *ku70Δ* and *lig4Δ*, are not sensitive to these insults (Manolis et al., 2001).

### H3-G34R but not H3-G34V cells exhibit genomic instability

H3-G34R cells exhibit genomic instability (Yadav et al., 2017). Since H3-G34V cells exhibit sensitivity to γ-IR, we asked if genomic stability was also compromised in H3-G34V cells. We initially monitored the loss of a non-essential minichromosome Ch16 (Niwa, Matsumoto, Chikashige, & Yanagida, 1989) from H3-G34V cells. As expected, cells lacking the heterochromatin protein Swi6^HP1^ displayed high frequencies of Ch16 loss (8.5%, (Fig. 2e)(Allshire, Nimmo, Ekwall, Javerzat, & Cranston, 1995). H3-G34R cells also lose the minichromosome at a significantly elevated frequency (4.3%) (Yadav et al., 2017), whereas H3-G34V and *set2Δ* cells display low levels of chromosome loss (2.2% and 1.8%), similar to H3-WT cells (1.2%).

As an alternate assay to measure chromosome segregation defects, we quantified chromosomal DNA segregation in late anaphase cells (with a spindle length> 10 microns) (Fig. 2f) (Ekwall et al., 1995). Since fission yeast have just 3 chromosomes, defects in chromosome segregation (lagging chromosomes) can be scored by monitoring the presence of DAPI-stained material at sites other than the ends of the spindle in late anaphase cells. We found that 7.4% of H3-G34R cells exhibited chromosome mis-segregation. In contrast, G34V and *set2Δ* cells showed no or little chromosome mis-segregation respectively (0.5%, 1%) compared to wild type cells (0.5%) and *clr4Δ* cells that lack heterochromatin (17.5%). We conclude from these assays that the genome of H3-G34V cells is relatively stable when compared to H3-G34R cells.

### Generation of a panel of H3-G34 mutant strains and analysis of H3K36me state

The surprising differences in the behavior of H3-G34V and H3-G34R mutants prompted us to ask whether H3-G34R phenotypes are caused by the large size or the basic charge of the arginine substitution, since valine is, like glycine, small and uncharged. To address this question, we generated a panel of sole copy H3-G34 mutant strains, with G34 replaced by the bulky uncharged residue tryptophan (W), the negatively charged glutamine (Q), the positively charged lysine (K) and Methionine (M) (Fig. 3a). Using the approaches outlined previously to determine whether antibodies against H3K36 methylation can still bind mutant H3 tails (Fig. 3b), we then assessed levels of H3K36me2 and me3 by western analysis on chromatin extracted from the H3-G34 mutant strains, and plotted quantification from western blots relative to H3-WT (Fig. 3c). This analysis showed that H3K36me3 is specifically reduced in H3-G34K (as well as H3-G34R and H3-G34V), but is not reduced in H3-G34W, M or Q mutants. Further, H3K36me2 is elevated in all H3-G34 mutant strains compared to H3-WT, when the reduced binding of anti-H3K36me2 antibodies to G34 mutant tails is taken into account.

**Figure 3.**
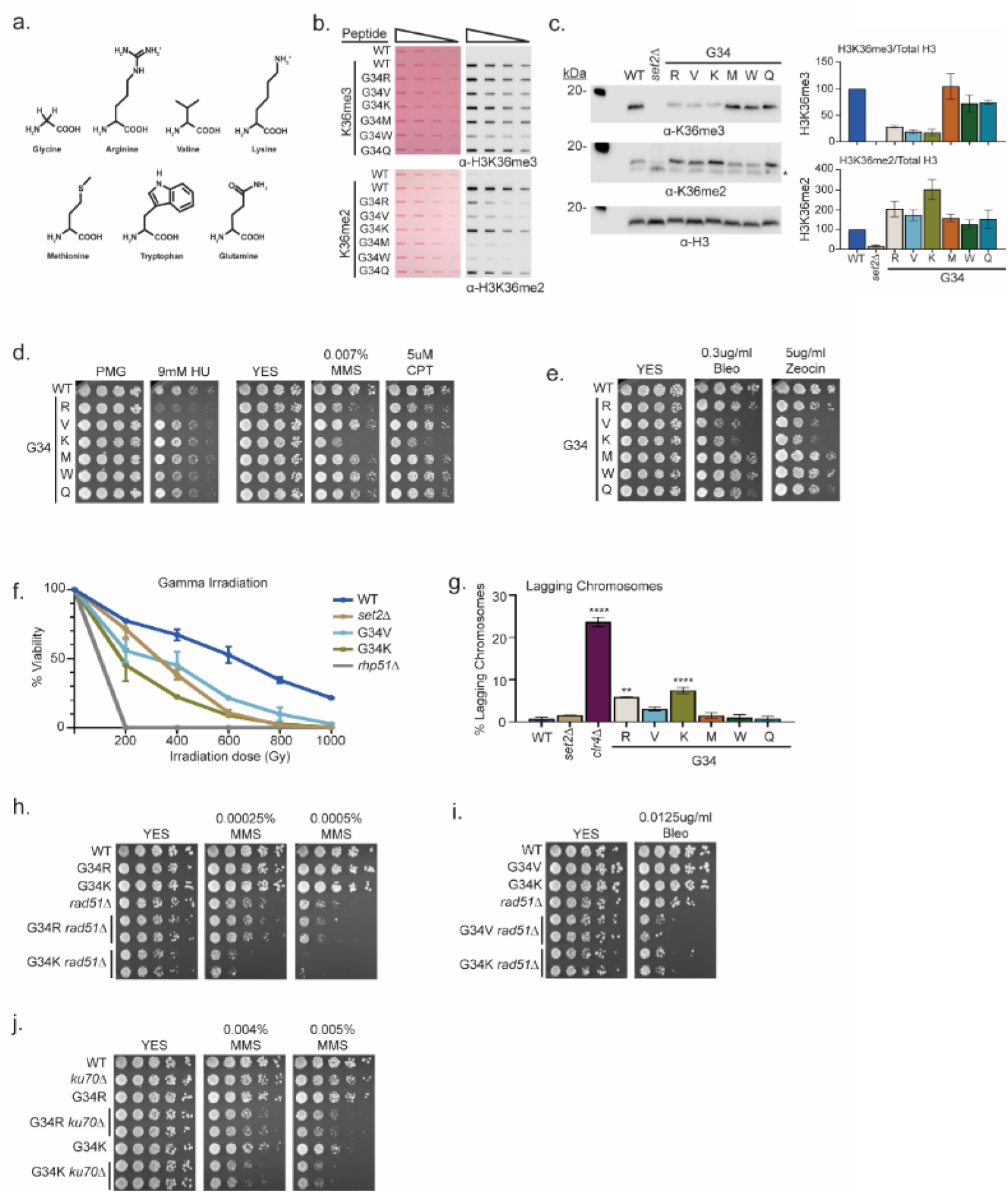
A panel of mutants at H3-G34 exhibit distinct effects on H3K36 methylation and different DNA-damage sensitivities. (a) Structure of glycine and amino acid substitutions used in experiments. (b) Dot blot analysis to quantitatively assess recognition of WT, H3-G34R, V, K, M, W, or Q peptides bearing K36me2 or K36me3 modifications by anti-K36 methyl antibodies. Peptides were loaded in 2-fold serial dilutions and ponceau staining was used as the loading control (left). (c) Western blot analysis of H3K36me3, K36me2, and total H3 in *set2Δ*, H3-WT, H3-G34R, V, K, M, W, and Q chromatin fractionated cellular extracts (left). The * symbol represents a nonspecific band in the H3K36me2 western. Quantification of K36 methylation levels relative to total H3 were calculated from 2 biological replicates (right). (d) Serial dilution yeast growth assay showing the effect of hydroxyurea (HU), methyl methanesulfonate (MMS) and camptothecin (CPT) on the indicated strains. (e) Serial dilution growth assay showing the effect of bleomycin and zeocin, two irradiation mimetics, on the indicated strains. (f) Effect of γ-irradiation (IR) exposure on viability of H3-WT, H3-G34R, H3-G34V, H3-G34K, *set2Δ* and *rad51Δ* cells. Data represent mean +/− SEM from 8 biological replicates. (g) IF analysis of lagging chromosomes in the indicated strains from 3 independent experiments. % lagging chromosomes represents the percentage of lagging chromosomes in anaphase cells counted. >200 anaphase cells were counted for each strain. ** represents significance difference of p<0.001 and **** a significance difference of p<0.0001 with H3-WT strain. (h) Serial dilution growth assay testing epistasis of H3G34R and H3G34K with HR pathway mutant *rad51Δ* cells. (i) Serial dilution growth assay testing epistasis of H3-G34V with *rad51Δ* HR-deficient cells. (j) Serial dilution growth assay testing epistasis of H3-G34R and H3-G34K with *ku70Δ* NHEJ-deficient cells.

### H3-G34 mutants show a range of DNA damage sensitivities and genomic stability

H3-G34K, V and R mutants showed loss of K36me3 and gain of K36me2, whereas H3-G34M, W and Q mutants retained H3K36me3 and gained K36me2. To ask the functional consequence of these changes, we assessed the sensitivity of the different mutant strains to DNA damage. We found that of all the H3-G34 mutants, only H3-G34R was sensitive to hydroxyurea. In contrast, both strains that bear the basic substitutions (H3-G34R and H3-G34K) were sensitive to the alkylating agent MMS and to topoisomerase inhibition CPT. We found that MMS sensitivity was not altered in strains bearing the bulky H3-G34W, G34M or the negatively charged G34Q substitutions (Fig. 3d). This would suggest that alkylation sensitivity is not linked to the size of the substitution at H3G34. On testing the γ-IR mimetics zeocin and bleomycin, we found that H3-G34K cells, like H3-G34V mutants showed sensitivity, whereas H3-G34M, G34W, G34Q and G34R cells did not (Fig 3e). We also monitored cell viability following γ-irradiation and found that H3-G34K and H3-G34V but not H3-G34R strains were sensitive to γ-IR, and that H3-G34K was more sensitive than either *set2Δ* or H3-G34V mutant cells (Fig. 3f). These experiments demonstrate the surprising complexity of DNA damage sensitivities of the different H3-G34 mutants, with some mutants (H3-G34V and H3-G34R) showing completely non-overlapping sensitivities, and others (H3-G34K and H3-G34R, and H3-G34K and H3-G34V) showing overlap for sensitivity to alkylating agents and IR respectively. Since all three mutants H3-G34R, H3-G34V and H3-G34K have reduced K36me3 and enhanced DNA damage sensitivity, K36me3 levels likely serve a critical role in safeguarding the genome, but the sensitivity of the different mutants to specific DNA damaging insults must be influenced by additional factors.

### H3-G34R and G34K exhibit chromosome loss but only H3-G34R appears defective for HR-mediated DNA repair

Since H3-G34K mutants were sensitive to DNA damage, we tested whether they showed genomic instability in the absence of exogenous DNA damage. Monitoring chromosome missegregation during anaphase, we found that H3-G34K cells had elevated levels of chromosome missegregation (7.4%) which were reproducibly higher than seen in H3-G34R cells (5.9%) (Fig. 3g). In contrast, none of the other H3-G34 mutants (V,M,W,Q) showed significant chromosome loss compared to H3-WT (0.7%).

H3-G34R cells are defective for HR-mediated DNA repair, and we previously proposed that chromosome missegregation in H3-G34R mutants might be due to the persistence of DNA repair intermediates in mitotic cells (Yadav et al., 2017). We therefore asked whether the other mutant exhibiting chromosome loss, H3-G34K, was similarly defective for HR-mediated repair. By introducing the *rad51* mutant allele and monitoring epistasis between H3-G34K and *rad51Δ*, we found that double mutant H3-G34K *rad51Δ* cells were more sensitive to both MMS treatment and bleomycin treatment than either single mutant alone (Fig. 3h and 3i), suggesting that H3-G34K is competent for HR. Similar results were found for H3-G34V (Fig. 2d and 3i). In contrast H3-G34R *rad51Δ* double mutants were no more sensitive to MMS treatment than *rad51Δ* alone (Fig. 3h), supporting that H3-G34R cells are defective in HR.

We also tested whether H3-G34K and H3-G34R mutants were deficient in NHEJ. H3-G34R or H3-G34K double mutants with *ku70Δ* showed elevated sensitivity to MMS compared with the single mutants alone (Fig. 3j), suggestive that NHEJ is intact in both the H3-G34R and H3-G34K strains. Unfortunately we could not perform this test for H3-G34V strains as they do not share genotoxin sensitivity with mutants defective for NHEJ. Together, these results suggest that HR is deficient only in H3-G34R mutants, and that NHEJ is intact in both H3-G34R and H3-G34K cells.

### H3-G34V, H3-G34R and H3G34K mutants show distinct transcriptional profiles

To try to rationalize the distinct phenotypes of H3-G34R, H3-G34V and H3-G34K mutants, we hypothesized that the mutations result in distinct transcriptional regulation. We performed RNA-seq in triplicate to compare transcriptional profiles of H3-G34R, H3-G34K, H3-G34V, *set2Δ* and H3-WT cells within the same experiment (Supplementary Table S4). Quantification of spike-in controls demonstrated that there was no global deregulation of transcripts in the mutant backgrounds, consistent with our previous data for H3-G34R and *set2Δ* mutants (Yadav et al., 2017). In total, 325 genes are upregulated, and 153 down regulated in *set2Δ* (employing cutoff +/− 1.5 fold, FDR 5%, (similar to the results we obtained previously (Yadav et al., 2017) and as initially reported in (3xH3) *set2Δ* strains (Matsuda et al., 2015; Suzuki et al., 2016)). H3-G34R, H3-G34K and H3-G34V exhibited fewer changes in gene expression, with 78, 128 and 94 genes increased in expression, and 82, 95 and 89 reduced, respectively (Supplementary Table S4). Comparison of genes that were differentially regulated between H3-G34R, H3-G34K and H3-G34V showed little overlap, with only 44 genes misregulated in all 3 strains (Supplementary Table S4).

Previously, our chromosome wide analysis of transcripts had revealed that genes that lie within the sub-subtelomeric regions of chromosomes I and II (herein designated as 120 Kb regions at the ends of chromosomes I and II) called ST domains (Buchanan et al., 2009; Matsuda et al., 2015; Yadav et al., 2017) were repressed in H3-G34R cells, whereas these domains are upregulated in *set2Δ* cells (Yadav et al., 2017). In this study, we found that the transcriptional profile of ST regions in H3-G34V cells more closely resembles that of H3-WT cells than the repression observed in H3-G34R, or the increased expression in *set2Δ* cells (Fig. 4a,b). In contrast, H3-G34K showed a slight trend toward transcriptional repression of ST regions (in particular on the left end of chromosome II). Further analysis of gene sets that were differentially regulated revealed that 16 of the 31 genes that were downregulated in both G34R and G34K strains were located within ST domains of Chr I or II, in contrast to just 2 of the 29 genes upregulated in both strains (Supplemental Table S4). We further confirmed the repression of ST domain gene expression in both H3-G34R and H3-G34K cells by RT-PCR analysis of *fah1*^*+*^ and *grt1*^*+*^ transcripts compared with controls (Fig. 4c).

**Figure 4.**
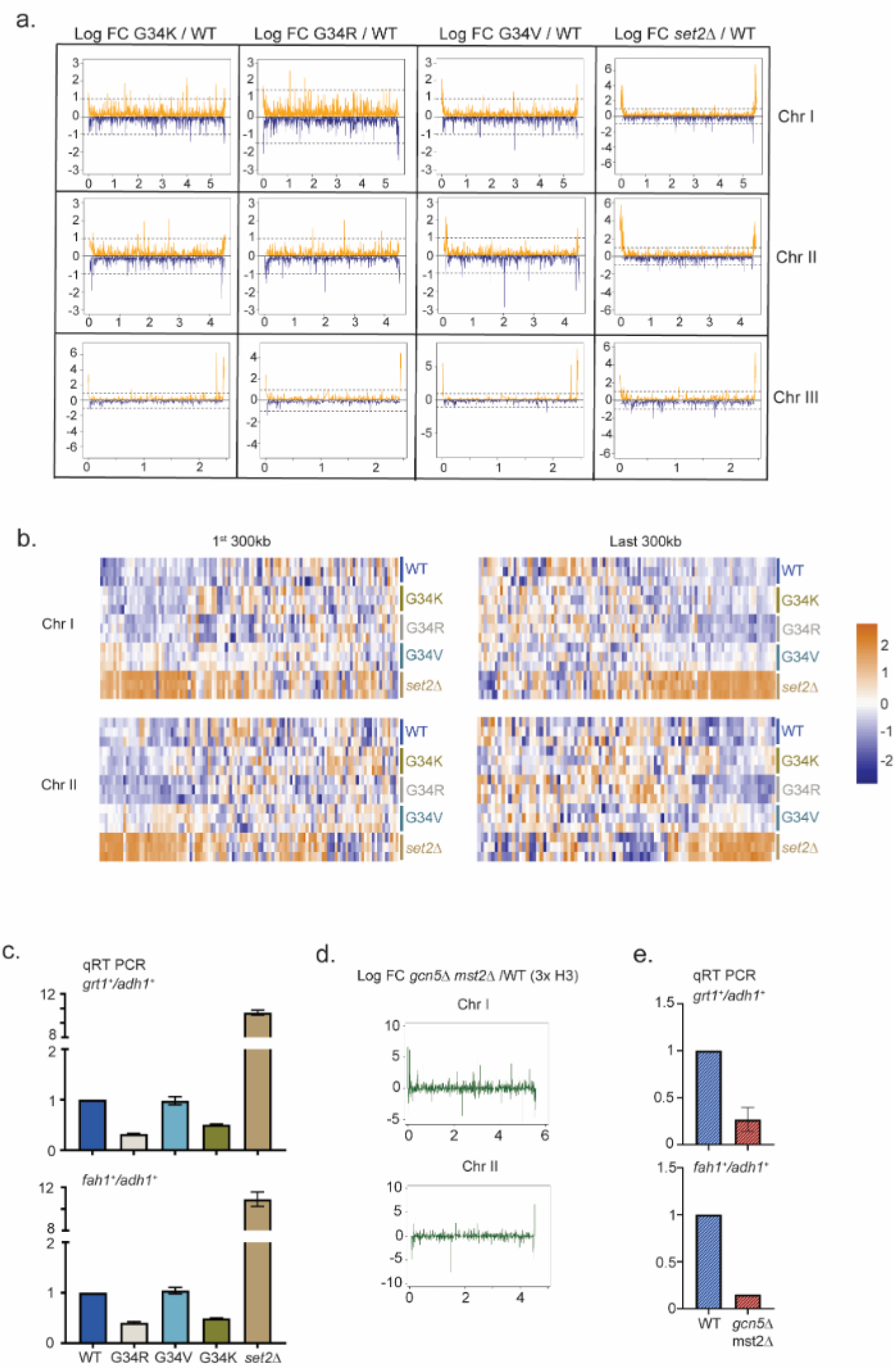
Distinct Transcriptional outcomes for H3-G34 mutants: with substitution with basic residues suppressing some subtelomeric transcripts, as seen in strains deficient in H3 K36 acetylation. (a) RNA-seq profiles for chromosomes I, II, and III comparing Log fold change ratios for H3-G34K/H3-WT, H3-G34V/H3-WT, H3-G34R/H3-WT or *set2Δ* /H3-WT plotted against chromosome coordinates. (b) Zoomed-in regions of Chr I (first 300Kb and last 300Kb, top) and Chr II (first 300Kb and last 300Kb, bottom) showing Z scores of log2 CPM for individual biological replicates. (c) qRT-PCR validation of ST genes *fah1*^*+*^ and *grt1*^*+*^ expression relative to *adh1*^*+*^ expression from 2 independent biological replicates. Samples were normalized to the WT-H3 strain. Subtelomeric transcripts in H3-G34R and H3-G34K are repressed compared with H3-WT and upregulated in *set2Δ*. (d) Chromosome-wide plots of transcriptional regulation in *gcn5Δmst2Δ* cells (3xH3) compared with wild type for Chr I and Chr II. Data reanalyzed from (Nugent et al., 2010). (e) qRT-PCR validation of *fah1*^*+*^ and *grt1*^*+*^ expression relative to *adh1*^*+*^ expression from 2 independent biological replicates. Samples were normalized to the WT-H3 strain. Subtelomeric transcripts in *gcn5Δmst2Δ* cells (3xH3) are reduced compared to wild type.

Intriguingly, the histone acetyltranferases Gcn5 and Mst2 also negatively regulate silencing of the ST domains at the termini of chromosomes I and II (Gomez, Espinosa, & Forsburg, 2005; Nugent et al., 2010) (Fig. 4d, data reanalyzed from (Nugent et al., 2010)), and by RT-PCR analysis of *fah1*^*+*^ and *grt1*^*+*^ expression we confirmed downregulation of ST gene expression in *gcn5Δmst2Δ* cells (Fig. 4e). Thus the downregulation of subtelomeric regions in G34R cells in which K36 acetylation is reduced is mirrored in cells lacking H3 K36 acetyltransferase function.

### H3-G34R cells and cells lacking H3K36ac accumulate Sgo2 at subtelomeres and have increased cytological ‘knobs’

Subtelomeric domains are surprisingly the most highly condensed regions of fission yeast chromatin and are microscopically visible as DAPI (4’,6-diamidino-2-phenylindole)-stained ‘knobs’ (Matsuda et al., 2015). Knob formation correlates with transcriptional silencing within subtelomeric domains, and requires Set2 H3K36 methyltransferase and the Shugoshin protein, Sgo2 (Matsuda et al., 2015; Tashiro et al., 2016). Sgo2 is an important component of the tension sensing machinery that assembles at centromeres during mitosis and relays turn-off of the mitotic checkpoint once kinetochores have properly attached to spindle microtubules (Watanabe, 2005). Recently however Sgo2 has been found at subtelomeres in interphase cells, where it promotes silencing, and regulates timing of replication of ST domains (Tashiro et al., 2016). Since H3-G34R but not H3-G34V shows extensive subtelomeric gene silencing, we examined “knob” formation in H3-WT, H3-G34R, H3-G34V and *set2Δ* cells. The levels of visibly condensed chromatin were similar in both single copy and (3xH3) wild type cells. Notably, the proportion of cells with visible knobs and their number per cell was significantly elevated in H3-G34R but not in H3-G34V cells (Fig. 5a), and as expected, knobs were reduced in cells lacking Set2 (Matsuda et al., 2015). A previous study showed that H3-K36Q mutant cells that mimic H3K36 acetylation and block K36 methylation dramatically lose knobs (Matsuda et al., 2015). Since Gcn5 acetylates H3K36, and subtelomeric domains are silenced in cells lacking Mst2 and Gcn5, we predicted that knobs would be increased in this mutant background. Indeed knob formation in *gcn5*Δ*mst2*Δ mutants is elevated, similar to levels in the K36ac-deficient H3-G34R cells (Fig. 5b).

**Figure 5.**
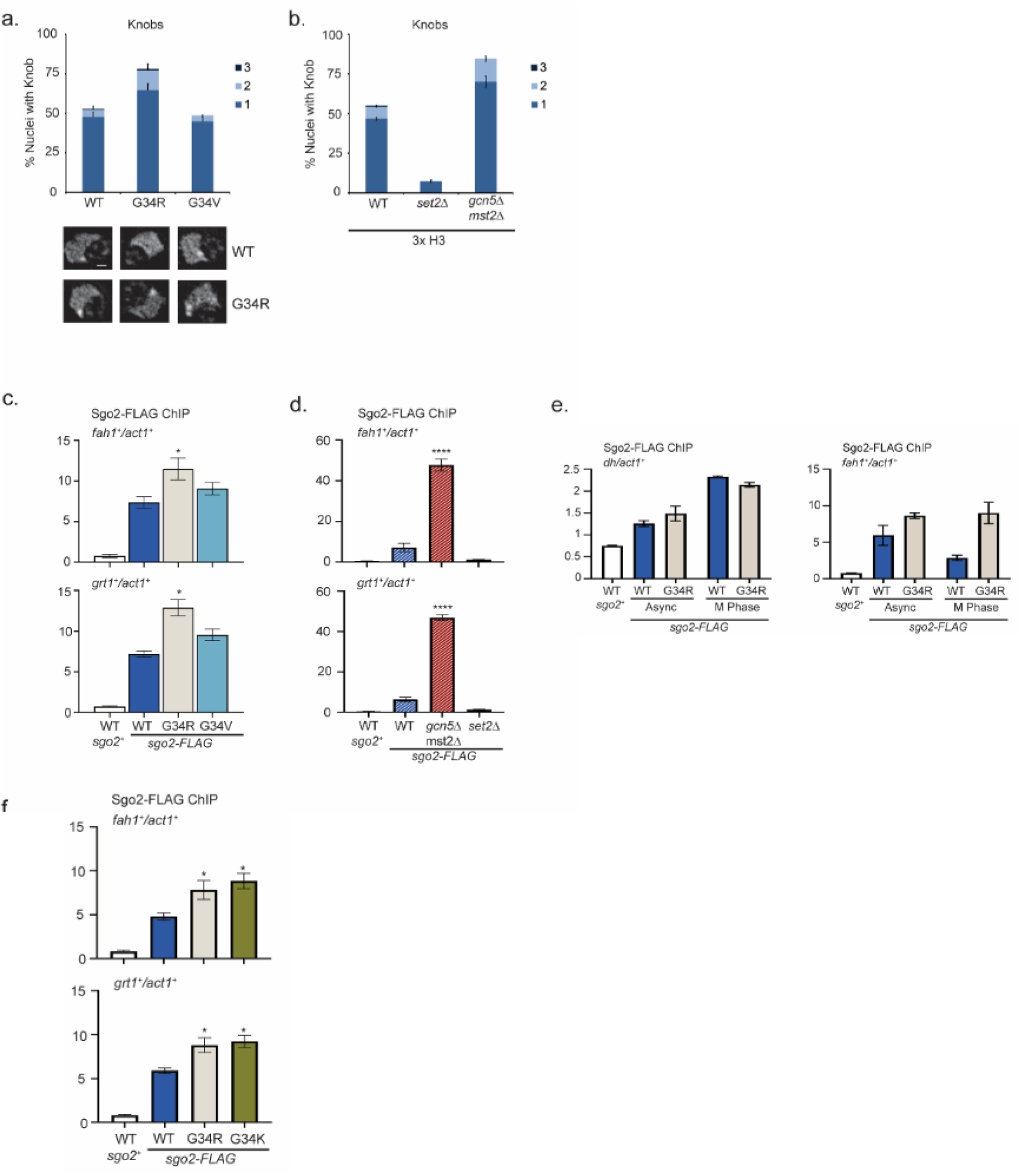
Formation of Knobs of highly condensed chromatin is enhanced in G34R and K36 acetyltransferase mutants, correlating with enhanced recruitment of Shugoshin to repressed subtelomeric domains. (a) Frequency of subtelomeric knob formation observed in H3-WT, H3-G34R, and H3-G34V cells. The number of knobs in a nucleus is shown. Data plots represent mean +/− SEM from 3 independent biological replicates, counting ~ 200 total nuclei per strain (upper). Example of nuclear knobs observed in WT and H3-G34; scale bar indicates 0.5 μm (bottom). (b) Frequency of subtelomeric knob formation observed in WT, *set2Δ*, and *gcn5Δmst2Δ* cells (all 3x H3) with the number of knobs in a nucleus shown. Data plots represent mean +/− SEM from 3 independent biological replicates, counting ~ 200 total nuclei per strain. (c-f) ChIP analysis of Sgo2-FLAG association with the subtelomeric *fah1*^*+*^ and *grt1*^*+*^ loci in (c) H3-WT, H3-G34R, and H3-G34V cells normalized to *act1*^*+*^. Data was collected and plotted as the mean of 6 biological replicates +/− SEM. * Represents a significance of p<0.05 compared with the H3-WT strain. (d) Sgo2-FLAG ChIP in (3xH3) WT, *set2Δ,* and *gcn5Δmst2Δ* cells normalized to *act1*^*+*^. Data is the mean of 4 biological replicates +/− SEM. **** Represents a significance of p<0.0001 compared with the WT strain. (e-f) Sgo2-FLAG ChIP at the subtelomeric *fah1*^*+*^ gene (e) or centromeric *dh* sequences (f) in asynchronous or mitotically arrested *nda3* mutant cells with sole copy H3-WT or H3-G34R (# replicates), and (g) Sgo2-FLAG ChIP in H3-G34K cells normalized to *act1*^*+*^. Data represents the mean of 6 biological replicates +/− SEM. * Represents a significance of p<0.05 compared with the H3-WT strain.

Since knob formation is dependent on Sgo2 (Tashiro et al., 2016), we hypothesized that Sgo2 may accumulate at ST domains in H3-G34R but not H3-G34V cells, and may also be enriched at ST in cells with reduced H3K36ac. Indeed ChIP revealed Sgo2-FLAG to be significantly enriched at ST domains in H3-G34R cells compared with H3-G34V or H3-WT (Fig. 5c), and that Sgo2 is highly enriched at ST domains in *mst2Δgcn5Δ* relative to WT controls (Fig. 5d). Since Sgo2 normally accumulates at centromeres in mitosis and is recruited to ST in interphase (Kawashima, Yamagishi, Honda, Ishiguro, & Watanabe, 2010; Matsuda et al., 2015; Tashiro et al., 2016), we asked whether H3-G34R influences the cell cycle dependence of its localization. Importantly, Sgo2 accumulation at centromeres in mitosis was unaffected by H3-G34R mutation (Fig. 5e), but even in mitotically arrested cells, Sgo2 was preferentially enriched at ST domains in H3-G34R cells compared to H3-WT (Fig. 5f), suggestive that in H3-G34R cells, cell cycle regulation of Sgo2 accumulation at ST is perturbed. In summary, our data suggest that Sgo2 accumulates at subtelomeres in H3-G34R or H3K36ac-deficient cells, correlating with enhanced silencing of ST domains and knob formation in these cells. Further, Sgo2 accumulates at ST domains in mitotic H3-G34R as well as interphase cells, although levels of Sgo2 at centromeres in mitotic H3-G34R cells are similar to H3-WT.

We also performed Sgo2 ChIP in H3-G34K cells. Consistent with enhanced Sgo2 recruitment to silenced ST domains seen in H3K36ac acetyltransferase defective and H3-G34R strains, H3-G34K cells also had elevated Sgo2 at ST sites (Fig. 5g). However, whether Sgo2 accumulation at ST correlates with reduced H3K36ac is unclear. Mass spectroscopic studies of K36 acetylation are not feasible on H3G34K because of the added complexity in data analysis from introduction of an additional lysine residue in the H3 tail, and western blot analysis is non-informative since antibodies against H3K36ac cannot effectively detect K36ac on G34 mutant H3 tails (Fig. 5-supplementary fig. 1a).

### Select phenotypes of G34R and G34K mutants show dominant effects

An intriguing feature of mutant histone oncoproteins is how they exert their effects when expressed in cells alongside higher levels of wild type H3 proteins. To determine which, if any, of the H3-G34 mutant phenotypes identified in our studies act dominantly, we generated “mixed H3” strains that express an H3-G34 mutant gene (*hht3.2*) and two wild type genes (*hht3.1*^*+*^, *hht3.3*^*+*^) from their endogenous loci (Fig. 1a), and performed comparative analyses with 3xH3 wild type strains.

We first asked if H3K36me3 was reduced in mixed H3 strains and found no change in H3K36me3 in mixed copy H3-G34R, H3-G34V or H3-G34K strains compared to 3xH3 WT (Fig. 6a). Similarly, we found no dominant increase in levels of H3K36me2 when the mutants were expressed in mixed H3 backgrounds (Fig. 6a). These results are consistent with the lack of dominant effect of G34R or G34V mutations on K36 methylation in mammalian cells (Lewis et al., 2013). Next we assessed H3 K36 acetylation. Due to the sequence difference at G34, we could distinguish K36ac peptides from both G34R and WT H3 proteins extracted from the mixed copy H3-G34R strain. We found that loss of H3K36ac was restricted to the G34R mutant H3 protein, whereas K36ac was present at comparable levels on wild type H3 purified from the mixed copy G34R and wild type 3xH3 strains (Fig. 6b).

**Figure 6.**
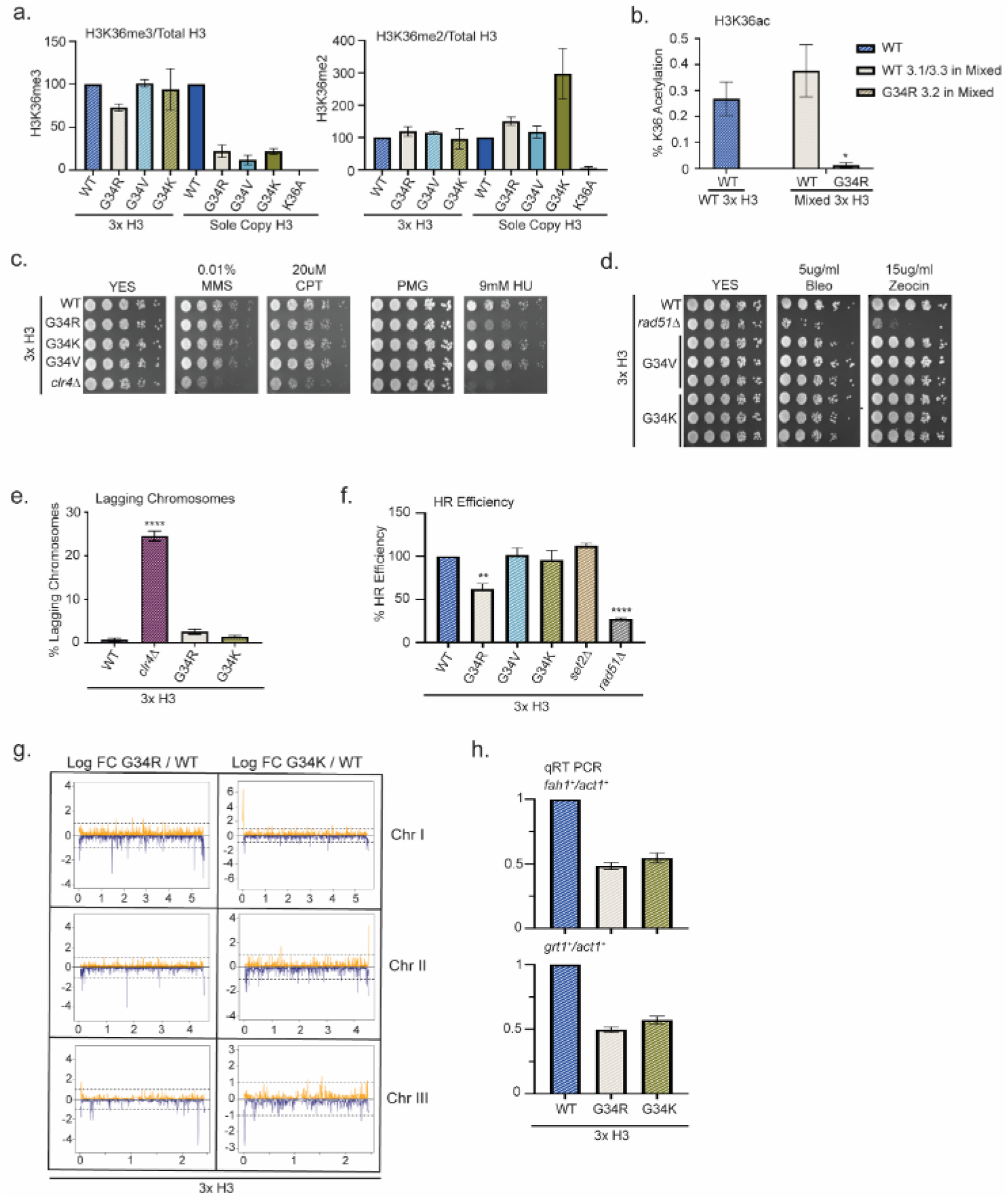
Dominance of HU sensitivity, HR defects and subtelomeric silencing in strains co-expressing H3-G34R and WT H3. (a) Western blot quantification of H3K36me2 and H3K36me3 normalized to total H3 in H3-WT, H3-G34R, H3-G34V, and H3-G34K chromatin extracts in both mixed H3 backgrounds (H3.1/H3.3 WT and H3.2 mutant) and sole copy H3 backgrounds. H3K36me2 and H3K36me3 methylation levels relative to total H3 were calculated from 2 biological replicates. (b) Mass spectrometry-based quantification of acetylation of H3K36 on histones purified from H3-WT (3x H3) and H3-G34R (mixed 3x H3 background) from 4 biological replicates. (c) 5-fold serial dilution growth assays showing the effect of HU, MMS, and CPT on the indicated 3xH3 strains. (d) Serial dilution growth assay showing the bleomycin and zeocin sensitivity of H3-WT, *rad51Δ,* H3-G34V, H3G34K, H3-G34V *rad51Δ* and H3-G34K *rad51Δ* 3xH3 cells. (e) Frequency of late anaphase cells that show a lagging chromosome in 3xH3 strains: WT, *clr4Δ*, H3-G34R, H3-G34K. Mean+/− SEM from 4-8 replicates. **** Represents a significance of p<0.0001 with the H3-WT strain. (f) Homologous recombination assay based on correction of leu1-32 mutation by HR as shown in Figure 2-figure supplement 1. Relative HR efficiency is shown as 100% for H3-WT (3xH3), and results are averaged from 3 independent experiments with error bars representing SEM. ** (p<0.005) and **** (p<0.0001) reflect significant differences with H3-WT cells. (g) RNA-seq profiles for chromosomes I, II, and III comparing Log fold change ratios for H3-G34R/H3-WT and H3-G34K/H3-WT plotted against chromosome coordinates. 3 biological replicates used for each strain, and all strains contain 3 copies of H3. (h) qRT-PCR validation of *fah1*^*+*^ and *grt1*^*+*^ expression relative to *adh1*^*+*^ expression from 2 independent biological replicates. Samples were normalized to the WT-H3 strain and shown as the mean +/− SEM. Subtelomeric transcripts in H3-G34R and H3G34K cells (3xH3) are reduced when compared with WT.

To ask whether DNA damage phenotypes were dominant, we assessed growth of 3xH3 strains on chronic exposure to the genotoxins MMS, CPT and HU. Mixed copy strains grew as well as 3xH3 WT strains, with the notable exception of H3-G34R (3xH3), which like the sole copy H3-G34R mutant, grew less well in the presence of HU (Fig. 6c, Fig. 6-supplementary fig. 1a). We also tested sensitivity to IR mimetics, and found that H3-G34K and H3-G34V mixed copy strains grew as well as WT (3xH3) strains on bleomycin and zeocin (Fig. 6d).

Our finding of the dominance of HU sensitivity of H3-G34R mixed copy strains led us to ask whether the H3-G34R mutation dominantly affected chromosome segregation or HR-mediated DNA repair. In contrast to the chromosome missegregation seen in H3-G34R and H3-G34K sole copy strains, there was no evidence of increased chromosome loss in the mixed H3 backgrounds for either H3-G34K or H3-G34R (Fig. 6e). There was however a significant drop in efficiency for H3-G34R mixed copy strains in our plasmid based HR DNA repair assay (Fig. 6f).

Finally we asked whether gene expression of H3-G34R and H3-G34K mixed copy strains differed from H3-WT (3xH3). We note that the overall numbers of genes that are differentially expressed in the (3xH3) background are reduced compared to the number of differentially regulated genes in the sole copy strains (54 genes differentially expressed in H3-G34R (3xH3) vs 160 genes in H3-G34R, and 142 genes differentially expressed in H3-G34K (3xH3) vs 223 in H3-G34K when using FDR 5% and FC of 1.5), suggestive that the presence of wild type H3 partially over-rides some transcriptional effects of the mutants. To determine whether the mutations exert dominant effects on expression of particular genes, we looked for overlap between genes that are differentially expressed in the single copy and mixed copy strains. Using the more relaxed criteria of p<0.05, we found that the majority of genes influenced by mutation in the (3xH3) background were also differentially regulated in the sole copy backgrounds (Fig. 6-supplementary fig. 1a, Supplementary Table S4). Manhattan plots indicated that subtelomeric domains of Chromosome I and II were repressed in H3-G34R mixed copy strains compared to WT H3 (3xH3), with a similar, but less marked trend also in H3-G34K mixed copy strains (Fig. 6g). Using real time PCR analysis of *grt1* and *fah1* transcripts, we validated the repression of subtelomeric genes in both H3-G34R and H3-G34K mixed background strains (Fig. 6h). Extending this analysis to ask if subtelomeric domains were preferentially repressed in H3-G34R or H3-G34K mixed copy strains (using FDR 5% and FC of 1.5), we found that a high proportion of genes repressed in G34R and G34K mixed copy strains were resident with ST domains (10/30 and 25/69 respectively) compared with only 1/24 and 7/73 of genes activated in G34R and G34K mixed copy strains (Supplemental Table S4). Thus many of the genes that are repressed by the H3-G34R or G34K mutation are situated in ST domains of chromosomes I and II, and the hyper-repression of gene expression within ST domains is dominant since it is evident in the mixed copy background strains.

## Discussion

In this work, we probe the consequence of mutation of Glycine 34 in histone H3 when the mutant provides the only source of H3, and identify phenotypes that dominate when mutants are co-expressed along with wild type H3. We found a surprising diversity of phenotypes with H3-G34 mutants differentially affecting post-translational modification of nearby K36, and DNA damage sensitivity and transcriptional control.

Given that a single enzyme, Set2, mediates all H3K36 methylation in fission yeast, it is intriguing that we only saw a reduction in H3K36me3, and only in some H3-G34 mutant backgrounds. Set2 may be unable to catalyze the di-methyl to tri-methyl transition on H3K36 when H3G34 is substituted by V, R or K. Consistent with a defect in tri-methylation, a very recent report shows that activity of the H3K36 tri-methylase SETD2 is blocked on nucleosomes bearing H3.3 G34 mutations, whereas NSD2-mediated di-methylation is unaffected (Jain et al., 2020). However, in that study, G34R, G34V and G34W similarly affected tri-methylation, whereas we saw no defect in H3K36me3 in the H3-G34W mutant, or H3-G34M or G34Q strains. This difference may stem from subtle differences in how H3 binds to the active site of SETD2 compared with Set2, resulting in differing outcomes for mutants such as H3-G34W in the 2 systems. Insight into Set2 and SETD2’s ability to accommodate G34 substituted H3 is limited because the only available structures are of SET2/D2 complexed with its high affinity ligand K36M H3.3 (Bilokapic & Halic, 2019; Yang et al., 2016; Zhang et al., 2017) which may influence the conformation of the active site. Another possibility is that the differences in K36me3 stem from our analysis of fission yeast H3 which differs from H3.3 at several residues including alanines instead of serines at residues 28 and 31 within the H3 tail (Matsumoto & Yanagida, 1985; Yadav et al., 2017), which may influence K36 methyltransferase behavior on G34 mutant templates (Schuhmacher et al., 2020). However, consistent with previous studies in mammalian cells, we found that coexpression of mutant (H3-G34R, V or K) with wild type H3 proteins did not affect H3K36me3, suggesting that loss of Set2-mediated trimethylation of K36 is limited to the mutant H3 tail.

K36 modifications have been linked to DNA damage response. In fission yeast, H3-G34R mutants with reduced K36ac and K36me3 are defective for HR-mediated repair of DNA breaks, whilst H3-G34V mutants, which maintain K36ac but have reduced K36me3, show no defect in HR. Since both mutants (and HR-proficient H3-G34K) have reduced H3K36me3, reduction in K36me3 is not causal of the HR defect. However, like H3-G34R mutants, Gcn5 mutants are reduced for H3K36ac, are HU sensitive and are also defective for HR (Pai et al., 2014), so these phenotypes may be linked. We confirmed that Gcn5 is the major H3K36 acetyltransferase in fission yeast and then showed in vitro that Gcn5 activity was lost specifically at K36 on recombinant G34R H3. This result emphasizes that H3-G34R directly affects Gcn5 function, but meant that we could not bypass the K36 acetylation defect in H3-G34R cells by Gcn5 overexpression to assign specific phenotypes to loss of H3K36ac. We also have not been able to use H3 mutants that mimic acetylated or non-acetylated K36 since they also impact K36 methylation. It therefore remains unclear whether the HR defect and sensitivity to HU seen in G34R cells are caused by loss of H3K36ac. These phenotypes show dominance in mixed copy H3-G34R strains, but there was no global loss of H3K36ac on wild type H3 purified from mixed copy H3-G34R strains. However, since chromatin is assembled from nucleosomes bearing 2 copies of H3, one possibility is that even in the presence of excess WT H3, H3K36ac is sufficiently diluted to cause dominant defects in HR and dominant HU sensitivity. In other words, the histone mutants disrupt the local chromatin architecture such that deposition of H3-G34R at *leu1* reduces local levels of K36ac and disrupts HR at that locus. Similar explanations have been suggested for local dominant roles of other histone mutants identified in cancer (Nacev et al., 2019). Unfortunately, we cannot readily determine if mixed copy (3xH3) G34R strains show global HR defects using *rad51Δ* in epistasis analyses since the mixed copy H3-G34R strain is only sensitive to HU at concentrations that kill *rad51Δ* in the chronic exposure plating assays. Additionally, we are unable to specifically track the localization of WT and mutant H3 proteins by ChIP since anti-H3G34R antibodies do not recognize fission yeast H3-G34R.

Another possibility would be that the unique phenotypes of H3-G34R mutants are due to the generation of a novel site of arginine methylation within the H3 tail. We have used mass spectroscopy to explore this possibility, but found no evidence for methylation of H3-G34R in fission yeast (Lowe, Maxham, Hamey, Wilkins, & Partridge, 2019).

In contrast to H3-G34R cells which are not sensitive to γ-irradiation, both H3-G34V and H3-G34K are sensitive to γ-IR and to IR-mimetics but there was no dominant effect since mixed copy strains were not sensitive to irradiation mimetics. This may mean that local perturbation of chromatin structure from deposition of the mutant histones is insufficient to yield irradiation sensitivity in the presence of wild type H3.

We also found that substitution of G34 with basic residues K or R led to accumulation of Sgo2 at subtelomeric (ST) domains correlating with enhanced silencing of these regions. These mutants also caused chromosome missegregation, but whether these phenotypes are linked is not clear. Cells defective for *gcn5Δ* and *mst2Δ* similarly show repression of ST domains (Nugent et al., 2010), and we found they accumulate Sgo2 at ST domains, and, like G34R, accumulate knobs. At ST domains, Sgo2 controls the timing of replication of late replicating origins, and loss of Sgo2 advances replication of some ST origins (Tashiro et al., 2016). In mitosis, Sgo2 is located at centromeres where it aids fidelity of chromosome segregation (Kawashima et al., 2010). One possibility is that the elevated levels of Sgo2 at ST domains in H3-G34R, H3-G34K and H3K36 acetyltransferase mutants further delays the timing of late-replicating ST origins, leading to chromosome mis-segregation, as well as replicative stress sensitivity. Alternately, the aberrant accumulation of Sgo2 at ST domains in mitotic H3-G34R cells could indirectly perturb centromere function. This is not due to depletion of Sgo2 from centromeres (as centromeric Sgo2 accumulation is similar between G34R and wild type cells), but perhaps from mislocalization and /or depletion of a limiting factor that is normally recruited by Sgo2 to centromeres in mitosis, and that in G34R cells is now retargeted preferentially to ST domains. However, we note that silencing of ST domains is maintained in mixed copy strains, whereas chromosome missegregation is corrected.

What is clear from many organisms is that recruitment of Shugoshin proteins relies on Bub1 kinase-mediated phosphorylation of histone H2A on S121 (T120 in mammals) (Kawashima et al., 2010; Kitajima, Hauf, Ohsugi, Yamamoto, & Watanabe, 2005; Kitajima, Kawashima, & Watanabe, 2004; Matsuda et al., 2015; Tang, Sun, Harley, Zou, & Yu, 2004; Tashiro et al., 2016). In fission yeast, both Bub1 kinase dead and H2A-S121A mutants lose localization of Sgo2 from centromeres and subtelomeres (Tashiro et al., 2016). We previously reported that H3-G34R mutants also alter the kinetics of phosphorylation at residues S128/129 of histone H2A (Yadav et al., 2017). Together with our new finding that H3-G34R/K mutants influence the localization of Sgo2, these data suggest the possibility that the G34R and G34K mutant H3s influence signaling through multiple phosphorylation sites clustered on the C-terminus of H2A. This is consistent with structural analyses of nucleosomes showing that the C-terminus of H2A lies close to the site of G34 substitutions on the H3 N-terminal tail (Du & Briggs, 2010).

We note that an interesting functional link between H3 and Shugoshin has also been made in budding yeast, where the homolog of Sgo2, SGO1 has been shown to bind histone H3. Although not reliant on G34 or K36 residues of H3, SGO1 binds a “tension sensing motif” at residues 42-45 on H3, which is necessary for Sgo1 recruitment to centromeres and for efficient chromosome segregation during mitosis (Deng & Kuo, 2018; Luo, Deng, Buehl, Xu, & Kuo, 2016). Intriguingly in this system, the histone acetyltransferase Gcn5 is important for ensuring efficient chromosome segregation when the tension sensing motif is damaged (Buehl et al., 2018; Luo et al., 2016). It is not clear whether Shugoshin proteins in higher organisms show a similar dependence on N-terminal regions of H3 for their localization and whether H3.3-G34R mutation influences their localization.

Finally we show dominant effects on transcriptional control exerted by the H3-G34R and G34K mutants. In particular, subtelomeric regions of chromosomes I and II are repressed. In keeping with the importance of transcriptional regulation in H3-G34 mutant cells, very recent work from the Lewis lab indicates that H3.3G34 mutants in mammalian cells exert profound transcriptional effects through modulation of Polycomb activity (Jain et al., 2020). They found that the decrease in K36me3 on H3.3 G34 mutant histone enhances PRC2 activity to promote elevated H3.3K27me3 in cis on the H3.3 tail (Jain et al., 2020). They propose that tumorigenesis driven by H3.3G34 mutations derives from enhanced Polycomb-mediated silencing of enhancers of genes that regulate differentiation, whereby cells retain a more stem cell like fate. Since acetylation at enhancers principally resides on K27 of histone H3.3 (Martire et al., 2019), this may explain why the G34 mutations identified in cancer occur solely on H3.3 (Behjati et al., 2013; Schwartzentruber et al., 2012).

H3.3 only differs from H3.2 by 4 residues: Ser31 in the tail, and three substitutions within the core of H3 which dictate chaperone selectivity (Ahmad & Henikoff, 2002; Elsasser et al., 2012; Liu et al., 2012). Phosphorylation of Ser 31 in H3.3 is important for regulation of cell fate during gastrulation and for enhancer activation (Martire et al., 2019; Sitbon, Boyarchuk, Dingli, Loew, & Almouzni, 2020) and Ser31P influences H3.3 K27 trimethylation and acetylation (Martire et al., 2019; Sitbon et al., 2020), and so phosphoregulation via S31 may provide an additional mechanistic link as to why the Gly 34 mutations are only found in H3.3.

Although fission yeast provides an exquisite system for analysis of histone mutations, we fully acknowledge the limitation of work in a system that lacks a Polycomb silencing pathway. It is interesting to note however the profound effects of G34R mutation on H3K36ac, the effects of the basic substitutions in the N-terminal tail of H3 on Shugoshin localization and the distinct DNA damage phenotypes of the different H3G34 mutations. It will be interesting to determine the extent to which the phenotypes uncovered in this study are shared in mammalian cancer cells expressing these mutants.

## Materials and Methods

### Strain generation

Histone mutant strains were generated as described (Mellone et al., 2003). Gene mutagenesis used standard PCR-based procedures, and strains are listed in Supplementary Table S1. All crosses used random spore analysis with nutritional/ drug/ temperature selection and PCR verification (including verification of loss of additional H3/H4 genes) and sequencing of H3.2 allele. Two independent clones for each genotype were used in nearly all experiments, and all experiments were performed at least twice.

### Yeast growth media

Fission yeast were maintained on rich (YES), or *pombe* minimal with glutamate (PMG) media with appropriate supplements (Moreno, Klar, & Nurse, 1991). PMG is Edinburgh minimal medium with glutamate.

### Plasmid DNA and recombinant proteins

Codon optimized Gcn5 (*S. pombe*) with 1 × FLAG was cloned into pET28a in frame with N terminal 6 × HIS tag by GenScript to generate JP-2587.

Fission yeast Histone H3 (and mutant versions (G34R, G34V, G34K)) were codon optimized by use of codon utilization analyzer 2.0 for expression in *e. coli* and the cDNAs were synthesized by IDT and were cloned into pCDF duet (Novagen) in frame with an N terminal 6 x HIS tag for bacterial expression: vectors JP-2395 (WT), JP-2489 (G34R), JP-2490 (G34V), and JP-2902 (G34K).

Recombinant pombe histones were purified from *e. coli* by “The Histone Source” at Colorado State University following standard procedures for histone purification (Dyer et al., 2004).

### Histone purification for Mass spectrometric studies

Histones were purified from fission yeast following a previously described purification protocol with some modification (Sinha et al., 2010). A 150 mL culture was inoculated to a density of 1.4 x10^6^ cells/mL in 4X YES media and grown at 25°C to a density of 3.6 ×10^7^ cells/mL and harvested by spinning at 3,000 RPM for five minutes. Cells were washed with H_2_O containing 10mM sodium butyrate, followed by NIB buffer (250mM sucrose, 20mM HEPES pH 7.5, 60mM KCl, 15mM NaCl, 5mM MgCl_2_, 1mM CaCl_2_, 0.8% Triton X 100, 0.5mM spermine, 2.5mM spermidine, 10mM sodium butyrate, 1mM PMSF, and Sigma yeast protease inhibitor). The pellet was frozen on dry ice and stored at −80°C. For lysis, the pellet was resuspended in 3 mL of NIB and transferred to two 7 mL bead beater tube along with chilled acid washed glass beads. Samples were cooled on ice and bead beaten twice for 2 ^1^/_2_ minutes at max power with 5 minutes on ice in between. The sample was collected by “piggy backing” into a 50 mL Oak Ridge tube at 3,000 RPM at 4°C in a benchtop centrifuge. The sample was then pelleted by centrifuging at 13,000 RPM for 10 minutes at 4°C in a Beckman Avanti centrifuge J-30I centrifuge using a JA25.50 rotor. The pellet was washed in 15 mL of NIB before resuspension in 10 mL of 0.4N H_2_SO_4_, sonication for 45 seconds at max power and incubation for two hours on a rotating wheel at 4°C. The sample was pelleted by centrifuging at 13,000 RPM at 4°C for 10 minutes and the supernatant was transferred to a new tube along with 5 mL of 5% buffer G (5% guanidine HCl and 100mM potassium phosphate buffer pH 6.8) where the pH was adjusted to 6.8 using 5N KOH. 0.5 mL of Bio-Rex™ pre-equilibrated in 5% buffer G was added to the sample and incubated at room temperature with rotation overnight. The resin was then washed twice with 20 mL of 5% buffer G and incubated with 3 mL of 40% buffer G (40% guanidine HCl and 100mM potassium phosphate buffer pH 6.8) for one hour at room temperature to elute the bound protein. Buffer exchange and concentration was performed against 5% acetonitrile with 0.1% TFA to a final volume of 150 μl and the sample was stored at −80°C. Protein concentration was measured and 10 μg of sample was electrophoresed and stained with coomassie blue for quality control.

#### Acetylation mass spectrometric analyses: UPLC−MS/MS Analysis

Acid extracted histone samples were TCA precipitated, acetone washed, and prepared for mass spectrometry analysis as previously described (Kuo et al., 2014). A Waters (Milford, MA) Acquity H-class UPLC system coupled to a Thermo (Waltham, MA) TSQ Quantum Access triple-quadrupole (QqQ) mass spectrometer was used to quantify modified histones. Selected reaction monitoring was used to monitor the elution of the acetylated and propionylated tryptic peptides. Transitions were created to study acetylation of pombe H3 wild-type and mutants as well as the H4 tails. The detailed transitions for peptides of H3 that vary in sequence from xenopus are reported in Supplementary Table S3, the transitions for the xenopus peptides and pombe H3G34R have been previously reported (Kuo et al., 2014; Yadav et al., 2017).

#### QqQ MS Data Analysis

Each acetylated and/or propionylated peak was identified by retention time and specific transitions. The resulting peak integration was conducted using Xcalibur software (version 2.1, Thermo). The fraction of a specific peptide (Fp) is calculated as

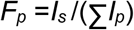

where I_**S**_ is the intensity of a specific peptide state and I_p_ is the intensity of any state of that peptide.

Data for acetylation analyses of H3-WT, H3-G34R and *set2Δ* came from 9 biological replicates, and 6 biological replicates for H3-G34V. 1-2 technical replicates from each prep were used, thus data for G34V was obtained from 9-12 samples, and for H3-WT, H3-G34R and *set2Δ* from 15-18 samples. Data to examine dominance using histones prepared from 3 X H3 strains used 6 biological replicates for 3xH3: WT, G34R, and *set2Δ*.

### Chromosome stability assays

#### 1. Minichromosome loss

The minichromosome (Ch16) (Niwa et al., 1989) bears an *ade6-216* allele which can complement the *ade6-210* allele present within the strain background. Loss of Ch16 causes loss of complementation of function of *ade6*^*+*^ and accumulation of red pigment when cells are grown on limiting adenine media. Strains containing Ch16 were grown in PMG –Leu (to maintain Ch16) at 32°C to a density of 5×10^6^ cells/mL. Cells were diluted in PMG (no additives) to a final concentration of 5-10×10^3^ cells/mL. Cells were plated onto PMG agar supplemented with amino acids and nucleobases with limiting (10% normal concentration) adenine, incubated at 25°C for 5 days and then transferred to 4°C to let red color develop. Ch16 loss frequencies were calculated by counting half sectored colonies (and those with >50% red but <100% red) and dividing by the total number of white, white sectored and less than half red colonies, excluding red colonies from the analysis as they have lost the minichromosome before plating. Results represent data from multiple (2-4) independent cultures of cells, and 5-10×10^3^ colonies were scored for each strain. Samples were assigned a code so identity was masked for this experiment.

#### 2. Lagging chromosome analysis

Chromosome mis-segregation frequencies were obtained as previously described (Mellone et al., 2003). >400 late anaphase cells were scored for presence of lagging chromosomes for each genotype, using at least 3 independent cultures of strains. Strain identities were masked for these experiments.

### Western and Dot Blots

#### Denatured extracts in 2X SB

Whole cell extracts (WCE) were made as published previously (Alper et al., 2013).

#### Chromatin fractionation

50 mL of cells were grown to a density of 6 x10^6^ cells/mL at 30°C in YES. Cells were collected by spinning at 3000 RPM for 3 minutes and washed 3 X with 1 mL of ice-cold NIB (described above). After washing, the supernatant was discarded, and the final chromatin pellet was resuspended in 1 mL of NIB. 500 uL of this material was mixed with 500 uL of 4X SDS sample buffer and heated at 98°C for 10 minutes. 10 – 20 uL was loaded onto a 12% SDS Page gel for analysis.

#### Antibodies used

**Anti-H3** (active motif pos, lot no. 26311003, Abcam ab1791). **Anti-H4** (millipore 05-858 lot no. 2020541). **Anti-FLAG M2** monoclonal (Sigma F1804), **anti-tubulin** (TAT1 kind gift from Keith Gull)(Woods et al., 1989). **Anti-H3K36me2**: Abcam ab9049, **Anti-H3K36me3**: Abcam ab9050, Cell signaling technology 4909. **Anti-H3K36Ac** (Abcam ab177179, rabbit monoclonal), Abnova (PAB31320), Rockland (600-401-I89), and Thermo Fischer (MA5-24672).

<u>Peptide sequences</u> used for assessment of K36 methyl Abs are listed in Supplementary Table S2.

<u>Slot blots</u> used a 2-fold serial dilution series with 150 uL of 50uM, 25uM, 12.5uM, 6.25uM, 3.125 uM, and 1.6 uM peptides spotted on activated PVDF 0.2 micron membrane using a 48 well BioRad Bio-Dot SF apparatus. Spots were air dried for 5 minutes before staining with Ponceau S stain to verify equal peptide loading. The membrane was blocked in 5% BSA in TBST at RT for 1 hour, incubated with primary Ab for 1 hour at room temperature, washed with TBST, incubated with HRP-conjugated anti-rabbit secondary Ab for 30 minutes, washed with TBST and then developed with enhanced chemiluminescence and images captured by LI-COR imaging. Anti-H3 K36 methylation Abs used are listed above.

#### Western Blot Quantification

All western blot quantification was done using LI-COR Image Studio software.

##### Chromatin immunoprecipitation

ChIP assays were performed similarly to (Alper et al., 2013), substituting Dynabeads Protein G (Invitrogen 1004D) for the protein G sepharose resin. Set2-FLAG ChIP used Anti-FLAG M2: (Sigma F1804) and monitored Set2 association with *act1*^*+*^ or *clr4*^*+*^ loci as a ratio of signal from input DNA; or Sgo2-FLAG association with subtelomeric regions, or centromeric regions relative to *act1*^*+.*^ Primer sequences used for q-PCR are listed in (Yadav et al., 2017). Sgo2 ChIP in the *nda3-KM311* background used asynchronous cells grown at 32°C or cells blocked in prometaphase by 8 hours incubation at 18°C.

##### Transcript analysis

###### RNA Seq studies

Hot phenol extraction was used to prepare the RNA (Leeds, Peltz, Jacobson, & Culbertson, 1991). 25 mL cultures were grown overnight in PMG complete media at 25°C to a density of 2.5 × 10^6^ cells/mL. The cells were pelleted by centrifugation and washed in DEPC H_2_O. The pellet was resuspended in 750 uL TES Buffer [50mM Tris-HCl pH 7.5, 10mM EDTA, 100mM NaCl, 0.5% SDS made in DEPC H_2_O] along with an equal volume of 5:1 phenol:chloroform pH 4.7 and incubated at 65°C for one hour with vortexing for 10 seconds every 10 minutes. The samples were then cooled on ice and centrifuged for five minutes at 13,000 RPM. The aqueous phase was transferred to a 2 mL phase lock tube and an additional phenol:chloroform extraction was performed. After centrifugation, the aqueous phase was transferred to a new tube and re-extracted using an equal volume of chloroform. To precipitate the RNA, the aqueous phase was transferred to a 2 mL microcentrifuge tube and three volumes of ice cold ethanol and 1/10^th^ volume 3M NaOAc pH 5.2 were added and the samples were kept at −20°C overnight. The next day, samples were centrifuged at 14,000 RPM and 4°C for 15 minutes to pellet the RNA. The pellet was washed with ice cold 70% ethanol and air dried for 30 minutes. A Turbo DNAse (Ambion) reaction was set up with 100 ug of RNA in a 150 uL reaction containing 5 uL of Turbo DNAse. The reaction was incubated at 37°C for 30 minutes and another 5 uL of Turbo DNAse was added and incubated for an additional 30 minutes prior to the removal of the DNAse using 50 uL of inactivation beads. An RNeasy Mini kit (Sigma) was used to further clean up and concentrate the RNA which was eluted in a final volume of 30 uL of DEPC H_2_O. 5 ug total RNA was diluted to 10 ul, and 2 ul of a 1:20 dilution of ERCC RNA Spike-in mix (Invitrogen) was added to each sample.

RNA-seq was performed by the St. Jude Hartwell Center. Ribosomal RNA was removed from the samples using Ribo-Zero Gold rRNA Removal Kit (Yeast) following manufacturer instructions (Illumina). RNA was quantified using a Quant-iT assay (Life Technology). The quality was checked by 2100 Bioanalyzer RNA 6000 Nano assay (Agilent) or LabChip RNA Pico Sensitivity assay (Perkin Elmer) before library generation. Libraries were prepared from 2 ug of RNA. Libraries were prepared using the TruSeq Strand Total RNA Library Prep Kit, beginning at Elution 2 – Fragment – Prime step immediately preceding cDNA synthesis according to the manufacturer instructions (Illumina) with the following modifications; the 94° C Elution 2 – Fragment – Prime incubation was reduced to five minutes and the PCR was reduced to 11 cycles. Libraries were quantified using the Quant-iT PicoGreen dsDNA assay (Life Technologies) or Kapa Library Quantification kit (Kapa Biosystems). One hundred cycle paired end sequencing was performed on an Illumina HiSeq 4000 (single copy strains) or on an Illumina Novaseq 6000 (triple copy strains). Three biological replicates were used for each strain analyzed. The total RNA was sequenced using stranded protocol with 2×100bp setting. The paired end reads were mapped to *S. pombe* (v2.29) genome using STAR (v2.5.3a) (Dobin et al., 2013). Reads counts for each gene were counted using the subread R package (Liao, Smyth, & Shi, 2019). Raw counts were TMM normalized and differentially expressed genes were analyzed using linear model of the mean-variance trend using the limma and voom packages in R. LogFC values were produced using the limma/voom packages (Law, Chen, Shi, & Smyth, 2014; Ritchie et al., 2015). External RNA spike-ins were analyzed to confirm that there were no changes in global RNA expression between the strain sets under analysis. Processed data is included in Supplemental Table S4. RNA-seq data has been submitted to GEO. Accession no. GSE # gse162572

**Real-time q-PCR** was performed on random primed cDNA generated from 2 independent RNA preps for each strain as previously described (Debeauchamp et al., 2008; Partridge et al., 2007) using primers that had been tested for linear amplification parameters, and working within the Ct range of linear amplification and using an Eppendorf Mastercycler® RealPlex^2^ machine. Transcript levels were normalized to *adh1*^*+*^ or *act1*^*+*^ transcripts.

### Observation of knobs

Knobs were observed as described previously (Matsuda et al., 2015). The *S. pombe* cells were cultured in liquid minimum medium with supplements (EMM2 5S) at 26°C with shaking to the early logarithmic phase. Cells were pelleted by gentle centrifugation, and chemically fixed by re-suspending in a buffer containing 4% formaldehyde (Polysciences, Inc., Warrington, USA), 80 mM HEPES-K, 35 mM HEPES-Na, 2 mM EDTA, 0.5 mM EGTA, 0.5 mM spermidine, 0.2 mM spermine and 15 mM 2-mercaptoethanol, pH 7.0. After fixation for 10 minutes at room temperature, cells were washed with PEMS (100 mM PIPES, 1 mM EGTA, 1 mM MgSO_4_, 1.2 M sorbitol pH 6.9) three times, then digested with 0.6 mg/mL zymolyase 100T (Seikagaku Biobusiness, Tokyo, Japan) in PEMS at 36°C for 5 minutes. Next, cells were treated with 0.1% triton X-100 in PEMS for 5 minutes, and washed three times thereafter with PEMS. Cells were incubated with 0.2 μg/mL DAPI in PEMS for 10 minutes, and then cells were resuspended with nPG-Glycerol (100 % glycerol for absorption metric-analysis (Wako, Osaka, Japan) with 4 % n-propyl gallate, pH 7.0) diluted with 1:1 with PEM (PEMS without sorbitol) and mounted on a clean 18×18 mm coverslip (No. 1S, Matsunami, Osaka, Japan). The slides were observed with 3D-SIM using a DeltaVision|OMX microscope version 2 or SR (Global Life Sciences Solutions Operations UK LTD) equipped with a 100x UPlanSApo NA1.35 silicone immersion objective or 60X UPlanApo NA1.42 oil immersion objective lens (Olympus, Tokyo, Japan). Reconstruction of 3D-SIM was performed by softWoRx (Global Life Sciences Solutions Operations) with Wiener filter constants of 0.003 using a homemade optical transfer function (OTF). Conspicuously condensed, DAPI-stained bodies were counted as knobs by visually inspecting each optical section of 3D-SIM images. Experiments were performed with three biological replicates for each strain, and a cumulative total of 189 to 293 nuclei were examined for each strain. Error bars represent the SEM.

#### Serial dilution analyses

Five-fold serial dilution assays were performed using exponentially growing cells and were spotted on agar plates using a plate replicator with 2 ×10^4^ cells in the 1^st^ spot. Plates were incubated at indicated temperatures. For chronic exposure assay, a fivefold serial dilution was spotted onto PMG complete agar plates +/− HU, on YES agar plates with DMSO or DMSO and CPT, and on YES agar plates +/− MMS. Plates were photographed after 4-5 days incubation at 30°C (CPT and MMS), or 6-7 days at 25°C (HU). All experiments were repeated at least twice.

##### γIR treatment

Cells at a density of 5×10^6^ cells/mL were irradiated using a Cobalt source, and 100 uL samples were taken following increments of 200 Gy exposure (Pai et al., 2014). Cells were plated on 6 YES plates for each assay condition, and colonies scored following incubation at 32°C for 4 days. The experiment was performed twice (4 biological replicates) to obtain average viability of treated versus untreated cells, and error bars represent the SEM. Strain identities were masked.

##### HR assay

We transformed *leu1-32* mutant cells (that bear a single nucleotide mutation in *leu1* that renders cells auxotrophic for leucine) with a fragment of wild type *leu1*^*+*^ (amplified using JPO 183/JPO 3480) and scored *leu1*^*+*^ transformants that can only arise by homologous recombination of the wild type *leu1* fragment into the mutant allele. The rates of HR were normalized by calculating transformation efficiencies of the different strains using *leu1*^*+*^ plasmid DNA (JP1050). The experiment was performed 3 – 4 X, with error bars representing SEM. Strain identities were masked.

### Statistical analyses

Statistical analyses were performed in GraphPad Prism version 7.03, using Ordinary one way Anova or Repeated measures Anova and Dunnett’s multiple comparisons test.

## Acknowledgements

We thank G. Zambetti for facilitating extension of RKY’s studies, L. Doorley and T. Hall for help with preliminary assays and R. Allshire, J. Tyler, S. White, S. Dent and M. O’Connell for useful discussions. JFP thanks L. Hendershot and T. Lawrence for support. We thank F. Bachand, S. Jia, E. Hidalgo, S. Forsburg, A. Annunziato, M. O’Connell, B. Strahl, R. Allshire and J. Kanoh for gifts of strains. We thank St. Jude protein production core for purification of Gcn5, St. Jude Hartwell Center staff for RNA-seq library preparation, DNA sequencing and peptide synthesis and J. Riggs (ARC) for help with the cesium source.

---

**Figure 1-figure supplement 1.**
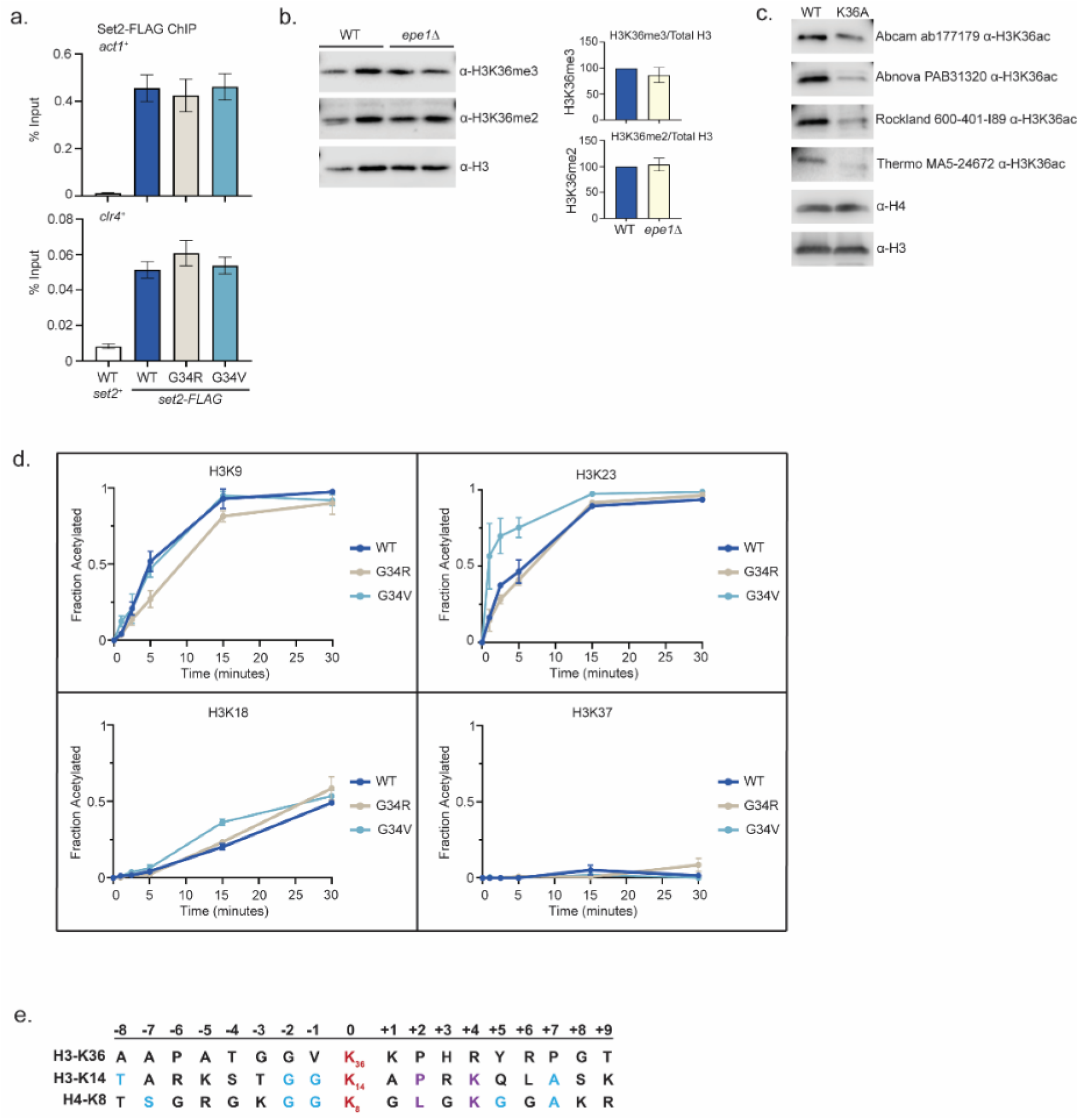
Differential modification of H3K36 in H3-G34R and H3-G34V mutants. (a) ChIP analysis of Set2-3xFLAG expressed from its endogenous locus in H3-WT, H3-G34V and H3-G34R cells. Set2-FLAG association with *act1*^*+*^ and *clr4*^*+*^ loci is represented as % of input DNA. Data represent mean +/− SEM from 6 biological replicates. No significant difference was found between WT and mutant H3 strains. (b) Western blot analysis of H3K36me3, H3K36me2, and total H3 in WT and *epe1Δ* cells using chromatin extracts. (right) Quantification of K36 methylation levels relative to total H3. (c) Western blot analysis of specificity of H3K36ac antibodies using WT and H3-K36A chromatin extracts (Abcam ab177179, Abnova PAB31320, Rockland Immunochemicals 600-401-I89, and Thermo Fischer MA5-24672). Total H4 and H3 were used as loading controls. (d) *In vitro* histone acetylation assay using recombinant Gcn5 and recombinant WT, G34R, or G34V H3, monitoring H3K9, H3K18, H3K23, and H3K37 acetylation. Data for each time point represents the mean +/− SEM from 3 biological replicates. K27ac was highly variable and was dropped from analysis. (e) Sequence flanking fission yeast H3K36 aligned with known and structurally analyzed GCN5 substrates H3K14, and H4K8. The sites of Gcn5 acetylation are shown in red while regions making protein contacts with GCN5 in previously characterized structures are denoted in purple. Aqua denotes residues conserved among known substrates. Alignment modified from (Poux & Marmorstein, 2003).

**Figure 2-figure supplement 1.**
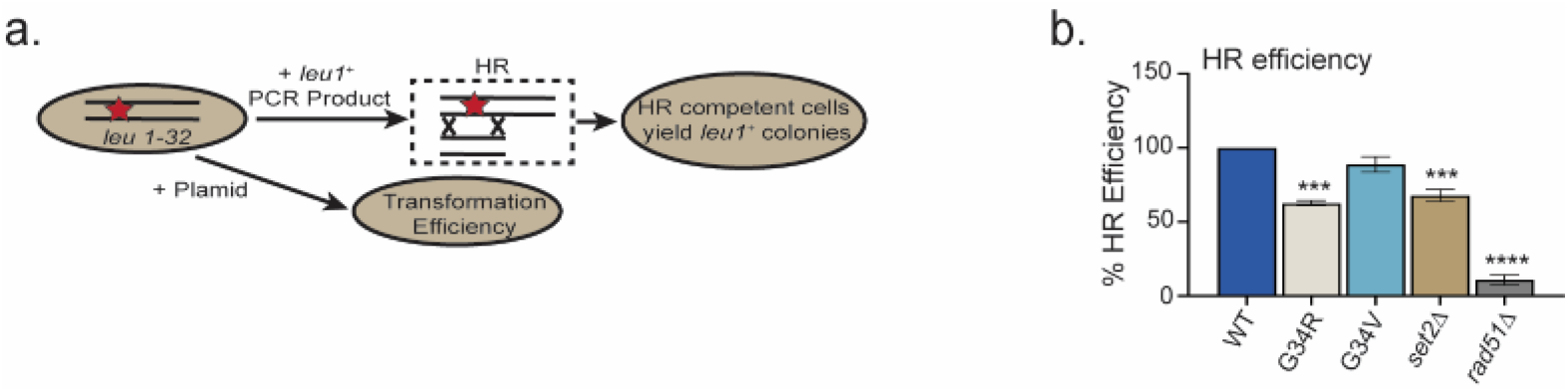
G34V exhibits no defect in HR. (a) Diagram of the homologous recombination assay based on correction of leu1-32 mutation by HR (Yadav et al., 2017). Cells of indicated genotypes were transformed with a *leu1* gene fragment to measure HR, or plasmid to measure transformation efficiency (top schematic). (b) Relative HR efficiency is shown as 100% for H3-WT, and results are averaged from 3 independent experiments with error bars representing +/− SEM. ** reflects significant differences with H3-WT cells ** (p<0.05), *** (p<0.001), and **** (p<0.0001).

**Figure 5-figure supplement 1.**
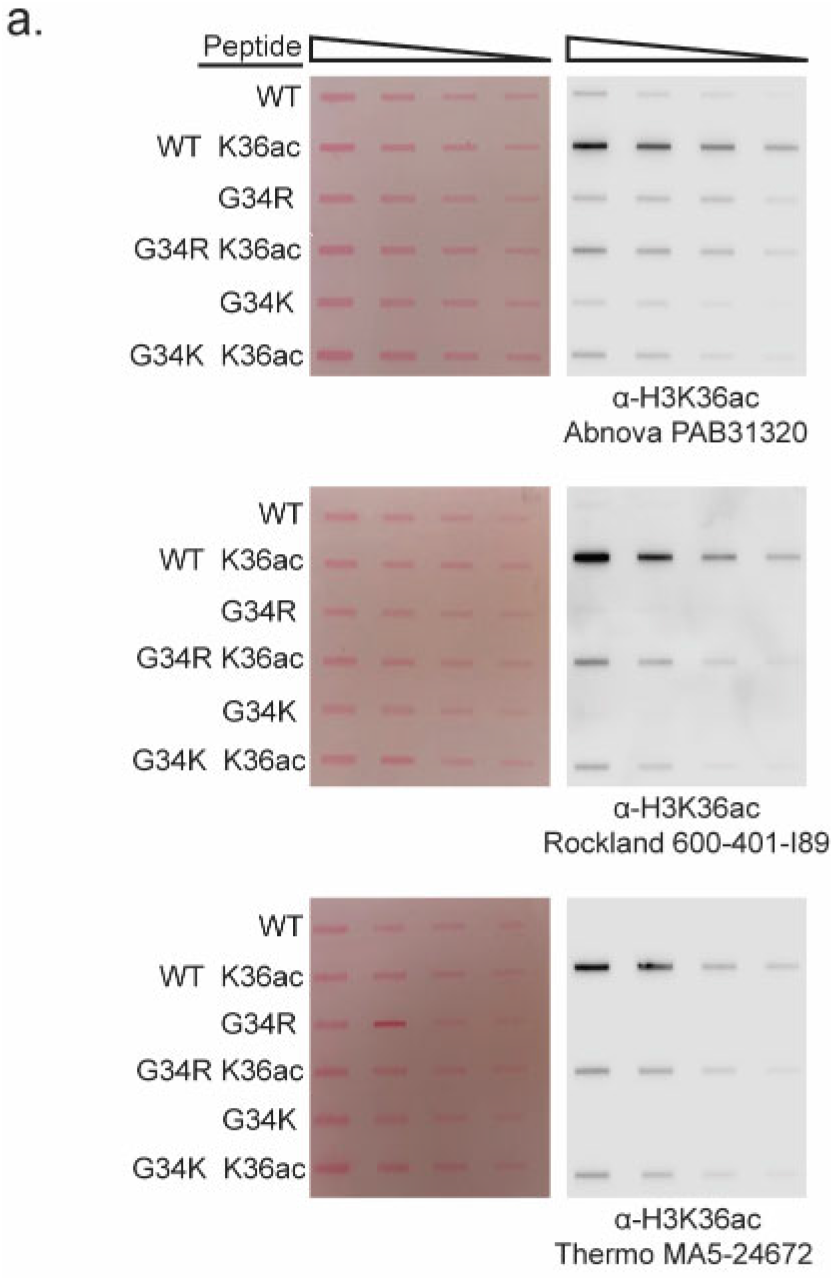
Determining effect of G34 substitution on antibody recognition of K36ac. (a) Slot blot analysis to quantitatively assess whether anti-K36 acetyl antibodies equivalently recognize WT, H3-G34R, or H3-G34K peptides bearing H3K36ac modification. Peptides were loaded in 2-fold serial dilution. Antibodies tested: Abnova PAB31320, Rockland Immunochemicals 600-401-I89, and Thermo Fischer MA5-24672. Ponceau stained blots were used as the loading control (left).

**Figure 6-figure supplement 1.**
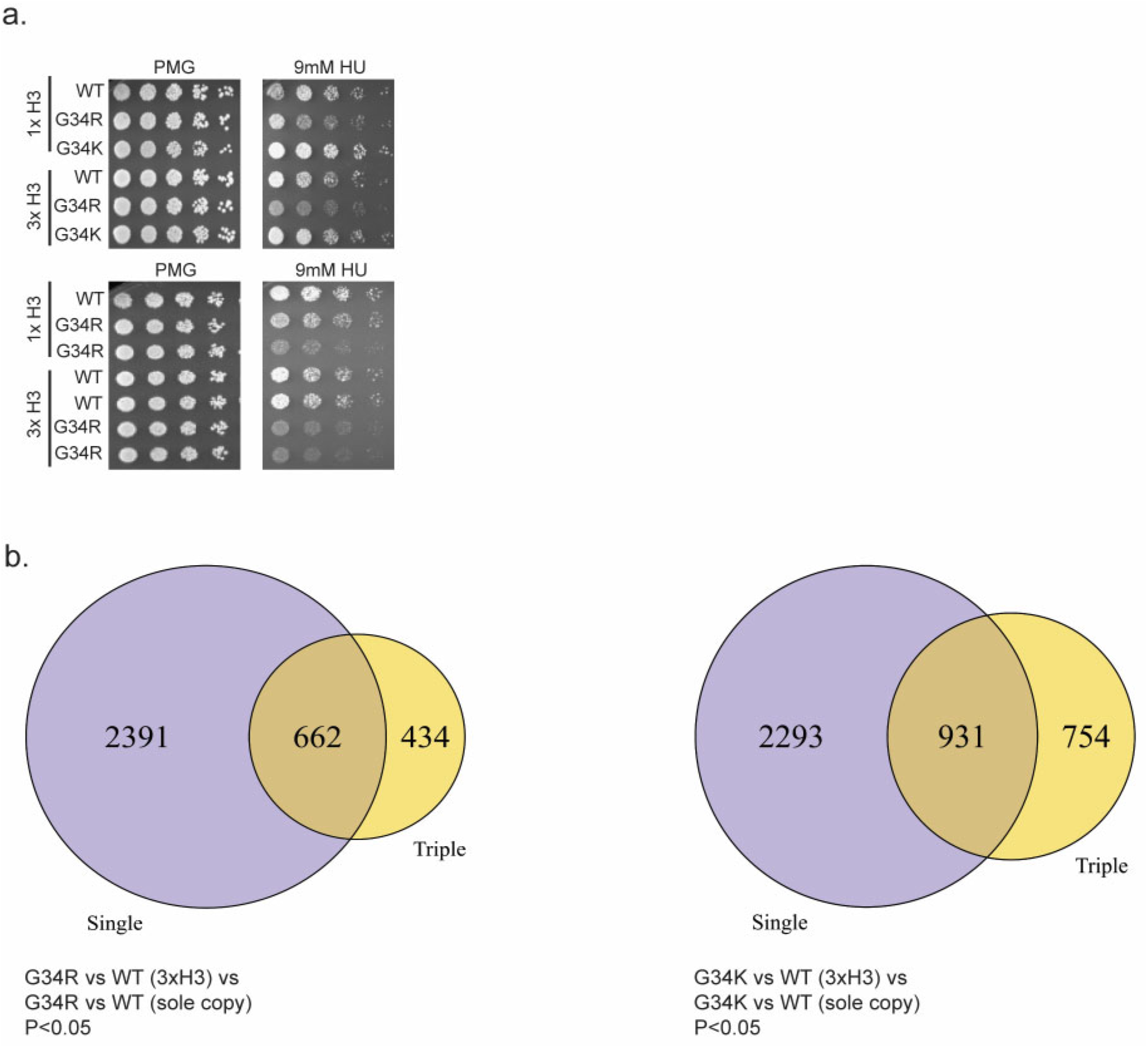
H3G34R exerts dominance over HU sensitivity, and both G34R and G34K dominantly suppress subtelomeric transcripts. (a) Replicate serial dilution growth assays to demonstrate HU sensitivity of G34R (3×H3) mixed background strain. (b) Overlap in gene expression profiles between H3-G34R sole copy and H3-G34R (3×H3) mixed copy strains relative to appropriate WT strains (left) and H3-G34K sole copy and H3-G34K (3xH3) mixed copy strains relative to WT strains (right) with cut offs of p <0.05 applied. The majority of gene expression changes seen in triple copy strains are in common with those in single copy strains.

We also include

**Supplementary Table S1**: Strain list.

**Supplementary Table S2**: Peptides used for Antibody characterization and for mass spectrometry calibration.

**Supplementary Table S3**: Detection parameters of unique tryptic peptides from *S. pombe* H3.

**Supplementary Table S4**: RNA-seq data.

